# Enabling Megascale Microbiome Analysis with DartUniFrac

**DOI:** 10.64898/2026.03.01.708916

**Authors:** Jianshu Zhao, Daniel McDonald, Igor Sfiligoi, Manuel E. Lladser, Lucas Patel, Yuhan Weng, Lora Khatib, Samuel Degregori, Antonio Gonzalez, Catherine A. Lozupone, Rob Knight

**Author notes:** Corresponding author: Rob Knight.

## Abstract

We introduce a new algorithm, DartUniFrac, and a near-optimal implementation with GPU acceleration, up to three orders of magnitude faster than the state of the art and scaling to millions of samples (pairwise) and billions of taxa. DartUniFrac connects UniFrac with weighted Jaccard similarity and exploits sketching algorithms for fast computation. We benchmark DartUniFrac against exact UniFrac implementations, demonstrating that DartUniFrac is statistically indistinguishable from them on real-world microbiome and metagenomic datasets.

## Main Text

UniFrac ^1-3^ is a phylogenetic beta-diversity metric that has been widely used in many microbiome and/or microbial ecology studies (>15,000), including large-scale ones such as Earth Microbiome Project (EMP) ^4^ and American Gut Project (AGP) ^5^, due to its capability to incorporate gene/genome evolutionary histories into community dissimilarity metrics. By leveraging branch-length information on the phylogeny, UniFrac frequently yields stronger between-group separation than non-phylogenetic distances (e.g., higher PERMANOVA R^2^ and clearer ordinations) such as Bray-Curtis and Jaccard dissimilarity, particularly when community turnover involves distantly related lineages, as shown in comparative evaluations across diverse datasets ^2^. As a fundamental biological community distance metric, UniFrac is not limited to amplicon sequencing-based community profiling techniques and it is natural to extend to metagenomic/genomic-based community profiling techniques as long as the unit of interest is clearly defined (e.g., ASVs/OTUs or metagenome assembled genomes (MAGs)-derived species unit based on Average Nucleotide Identity, or ANI) ^6^. The computational complexity of UniFrac (weighted) is proportional to the number of taxa in the phylogenetic tree and is quadratic in the number of samples in a given study (pairwise comparisons) ^7^. Model-based microbial diversity estimates with the 16S rRNA gene suggest that Earth may harbor more than 10^12^ microbial species ^8^, or even more with whole genome-based approaches. But we are far away from capturing these diverse microbial species. As high throughput sequencing is becoming readily accessible in standard microbiome studies, we can now sequence thousands of samples with millions of species in a single project. This, however, has created a major challenge—computing all-versus-all UniFrac distances becomes a bottleneck for real-world datasets with even more samples and taxa. Many computational optimizations have been developed for faster UniFrac computation for large numbers of samples (e.g., a few thousand) over the past 20 years ^7, 9, 10^, e.g., faster tree traversal strategies ^10^, better parallel efficiency (Striped UniFrac) ^7^ or hardware acceleration (e.g., Simple Instruction Multiple Data/SIMD and GPU) ^9^. However, these improvements were based on the same exact UniFrac algorithm and cannot scale further.

Here, we invented a new UniFrac algorithm, DartUniFrac, that can scale to millions of samples with billions of taxa. First, we revisited original UniFrac ^1^ (unweighted, Fig. S1a): 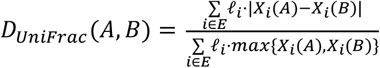, where 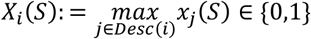, and *x*_*j*_(*S*) is defined as: 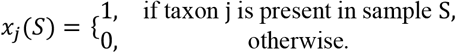. Weighted UniFrac is defined as: 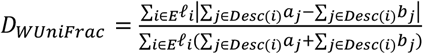 (Fig. S1b and Supplementary Materials, a different but equivalent equation to the original Weighted UniFrac) ^2^. We prove that unweighted and weighted UniFrac are essentially weighted Jaccard similarity on tree branches (Supplementary Methods and Fig. S1a and b). For unweighted, *D*_*UniFrac*_ =1 − *J*_*w*_ (*x,y*) where 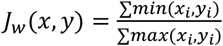 (also known as weighted Jaccard similarity), *x*_*i*_ = ℓ ∗ *X*_*i*_ (*A*), *y*_*i*_ = ℓ ∗ *X*_*i*_ (*B*) (Fig. S1a). For weighted, 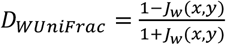, where 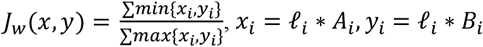 and 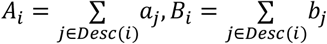 (Fig. S1b). In both unweighted and weighted UniFrac, computing weighted Jaccard similarity 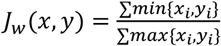 is the limiting step because for real-world datasets, the dimension of *x* and *y* (equal to the number of branches in the phylogenetic tree) can be millions or even billions. The problem becomes how to compute weighted Jaccard similarity fast and efficiently. First, we use an optimal balanced parentheses (BP) data structure to represent phylogenetic trees so that key navigation operations (e.g., find parent/child/sibling) in a tree run in constant time (Fig. 1a and Online Methods). This allows representation and efficient traversal of trees with billions of taxa. Each branch of the tree is now a set element, but the weight of each set element is related to the samples, either presence or absence of taxa that are descendants of the branch (unweighted) or sum of relative abundances of taxa that are descendants of the branch (weighted) (Fig. 1a, Fig. S1a and b and Supplementary Methods). After obtaining the weighted sets (Fig. 1a), computing weighted Jaccard similarity is the most expensive step because for real-world microbiome datasets, there can be billions of taxa/branches and millions of samples. We rely on weighted MinHash (also known as sketching algorithms) to sketch weighted sets for computing weighted Jaccard similarity. MinHash belongs to a category of locality-sensitive hashing algorithms that is widely used in data mining for large-scale web and text comparison. MinHash is also widely used in genomics and metagenomics for computing average nucleotide identity (or ANI) at large scale ^11-14^, and it allows for the computation of weighted Jaccard similarity efficiently with controllable estimation error, which converges to 0 as the sketch size increases (Fig. 2a and Online Methods). Among several weighted MinHash algorithms, DartMinHash and Efficient Rejection Sampling (ERS) are the most efficient for sparse and dense sets, respectively ^15, 16^. In the resulting sketch (a low-dimensional representation with length S of original weighted sets from all branches), computing an integer-based Hamming similarity (normalized) between each pair of sketches estimates weighted Jaccard (*J*_*w*_) and thus also unweighted UniFrac or weighted UniFrac (Fig. 1a and Online methods). Real-world microbiome datasets are sparse, such that most branch sets are empty, making the Weighted MinHash sketch step extremely fast with DartMinHash being the fastest weighted MinHash algorithm both in theory and in practice ^15^, to the best of our knowledge. Overall big-O notation for DartUniFrac is O(N*T_active_ + N*S*log(S) + N^2^*S), where N is the number of samples, T_active_ is the number of active branches on average across all samples, and S is the Weighted MinHash sketch vector length (See detailed big-O analysis in Online methods). Big-O notation for exact UniFrac is O(N^2^*T), where T is the total number of branches or taxa and in practice, S<<T (S is less than or equal to 2,048 in practice). The computational bottleneck for DartUniFrac for millions of samples is dominated by the pairwise integer Hamming similarity step (O(N^2^*S), which computes equal slots between sketch vectors with the same length, where the slots are 16-bit integers) (Fig. 1a and Online Methods). We first maximize CPU performance for this step using multithreading and SIMD. However, Hamming similarity computation is memory-bandwidth-bound, so further speedups on CPU are limited (e.g., more CPU threads), but can benefit from hardware with higher memory bandwidth. We therefore offload this step to GPUs to further speed up the computation for large numbers of samples (Fig. 1a). A streaming mode (both CPU and GPU implementation), which allows block-by-block computation of the full distance matrix is also provided when the distance matrix cannot fit into RAM (Fig. 1a).

**Figure 1.**
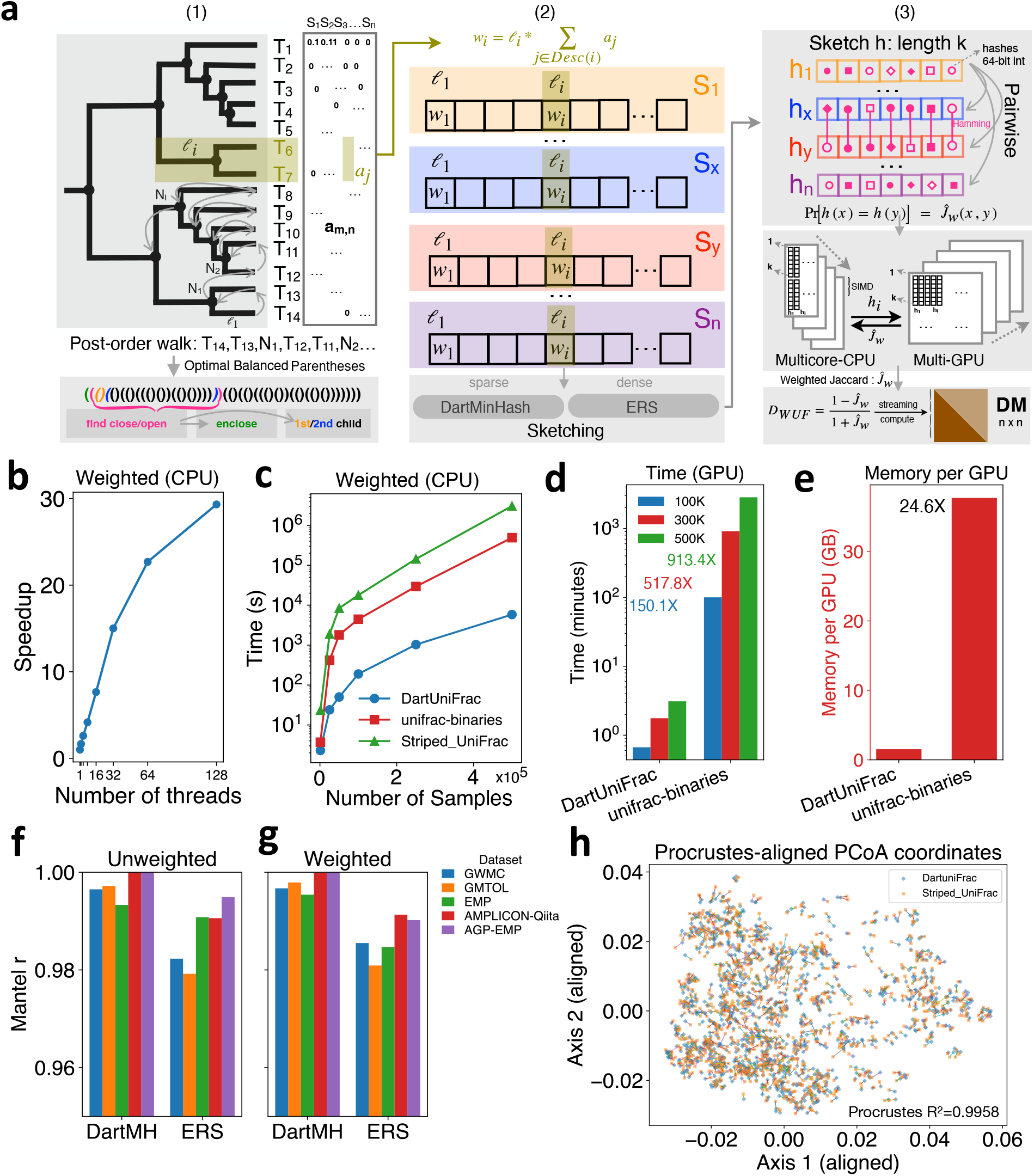
(a) Schematic overview of the DartUniFrac algorithm and implementation. The phylogenetic tree was represented by optimal balanced parentheses for efficient retrieval of branch length relevant for each sample. Then the input set vectors (each dimension represents a branch of the tree, skipped if not relevant for that sample, there can be millions of branches or more for big trees) were collected and passed to sketching algorithms, DartMinHash (default) or ERS. The output from sketching is a much smaller vector (1,000-2,000, called sketch vectors) storing 64-bit integer hashes (truncated to lower b bits afterwards, see Supplementary methods). The number of equal hashes for all slots in the sketching vector out of total slots equals weighted Jaccard similarity. CPU multi-threading and SIMD, or multi-GPU can be used to speedup pairwise computation of equal hashes. (b) Scalability of DartUniFrac CPU implementation with respect to the number of CPU threads. (c) Speedup of DartUniFrac-CPU over Striped UniFrac and unifrac-binaries-CPU (hardwared optimized Striped UniFrac, here SIMD for unifrac-binaries-CPU) for Weighted UniFrac with respect to the number of samples. (d) Speedup of DartUniFrac-GPU over unifrac-binaries-GPU for 100K, 300K and 500K samples. This benchmark was performed on Intel(R) Xeon(R) 6972P 96-thread CPU with NVIDIA A100 (HBM2e) GPUs. (e) GPU memory requirement for DartUniFrac-GPU and unifrac-binaries-GPU. Mantel correlation coefficient between distance matrices estimated by DartUniFrac and Striped UniFrac (truth) for both unweighted (f) and weighted UniFrac (g) across 5 datasets (see also Table S6). (h) Procrustes analysis between DartUniFrac and Striped UniFrac for the Global Water Microbiome Consortia (GWMC) dataset (unweighted). Procrustes R^2^ is shown in the plot. See Figure S7 for weighted.

**Figure 2.**
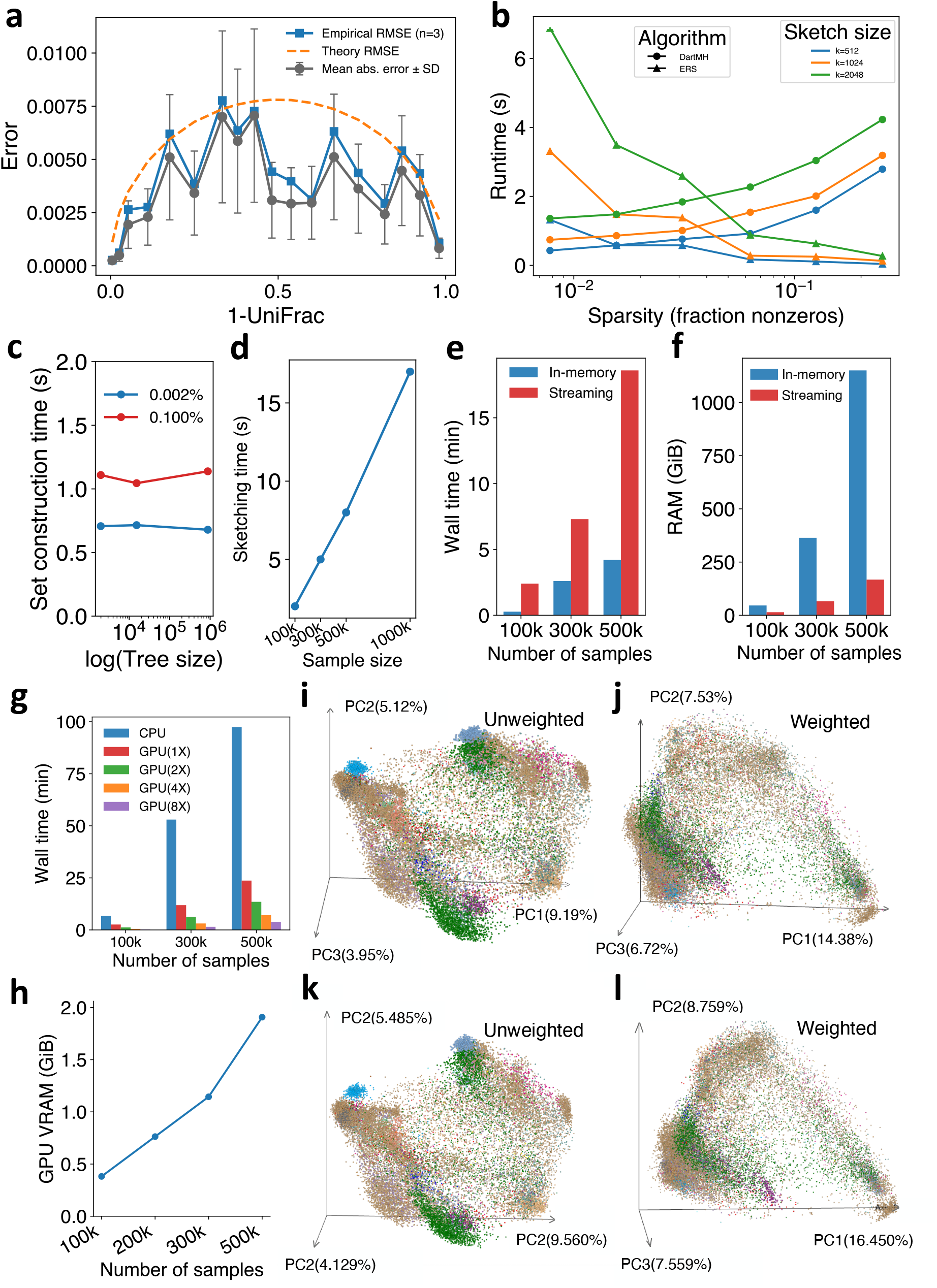
(a) Estimation error of our DartMinHash implementation for simulated UniFrac distance (0,1), including Rooted Mean Square Error (RMSE), absolute error and theoretical error. A sketch size 2048 was used for this analysis. (b) Running time versus sparsity of input weighted sets for DartMinHash and ERS. Sparsity was measured as fraction of nonzeros considering all the set/branch elements. (c) Branch set construction time with respect to number of taxa and sparsity while fixing number of samples. (d) DartMinHash sketching time with respect to number samples. (e) Running time comparisons between DartUniFrac-GPU in-memory mode and streaming mode. (f) Peak RAM comparisons between DartUniFrac-GPU in-memory mode and streaming mode. (g) GPU running time for DartUniFrac versus CPU running time for DartUniFrac with respect to the number of samples and number of GPUs. (h) (i)-(l) PCoA visualization for the Earth Microbiome Project (EMP) dataset for DartUniFrac (i and j, unweighted and weighted respectively) and Striped UniFrac (k and l, unweighted and weighted respectively, exact UniFrac). PCoA explanation rate by each principal coordinate was labeled next to each axis. Data points were group by environmental type. Detailed legend for (i), (j), (k) and (l) can be found in Figure S9.

DartUniFrac-CPU scales well with increasing number of CPU threads (Fig. 1b) and is more than 200 times faster than the state-of-the-art UniFrac algorithms (Striped UniFrac ^7^ and a hardware optimized version, called unifrac-binaries ^9^, both are exact UniFrac algorithms) for in-memory mode (Fig. 1c, Table S1). Additionally, DartUniFrac-CPU can compute pairwise UniFrac for a million samples (87,522 taxa) in 1.8 hours on CPU (Table S1) provided enough RAM is available, or around ∼4.5 hours in a streaming mode when there is not enough RAM (Table S1). Notably, the existing state-of-the-art unifrac-binaries-CPU required more than 20 days to complete the same task on a CPU. In streaming mode, memory requirements can be one order of magnitude smaller (Fig. S2) for DartUniFrac-CPU. GPU support provides another ∼20 times speedup than CPU (Table S2 and Fig. 2g). DartUniFrac-GPU is on average ∼900 times faster than the unifrac-binaries-GPU implementation (Fig. 1d) while consuming ∼24 times less GPU memory for ∼87,522 taxa and 500,000 samples (Fig. 1e). DartUniFrac-GPU can be even more GPU memory efficient as the number of samples further increases (Online methods). DartUniFrac allows computing UniFrac distances beyond the limitations of BIOM-Format version 2.1.0 ^17^, a common container for microbiome feature tables, which is capped at 2^32^ nonzero values in the feature table; with feature tables that extend the nonzero limitation to 2^64^, DartUniFrac-GPU can finish in 13.8 minutes for 500,000 samples with 20 million taxa using 2 GPUs (see detailed hardware information in Table S3). Because DartUniFrac-GPU only stores the sketch vector (16-bit per element) for each sample in GPU memory, it can easily scale to millions of samples with moderate GPU memory (e.g., 48G is enough for 10 million samples with default sketch size) (Fig. 2h). Integer Hamming similarity is memory-bandwidth-bound and we observed significant speedup on the most recent GPUs than on CPU (Fig. 2g, Table S2, S3, S4 and S5). In DartUniFrac-GPU streaming mode, it can be 2 times slower than in-memory GPU mode but requires ∼10 times or smaller CPU RAM (Fig. 2e and f). For 1,000,000 samples or more, DartUniFrac-GPU is > 1,000 times faster than unifrac-binaries-GPU in terms of compute time as both need the same IO time (Table S4).

PCoA analysis on EMP and GWMC (Global Water Microbiome Consortia) datasets ^18^ showed that DartUniFrac provides nearly identical results with exact UniFrac distances as measured by Mantel and Procrustes analysis on the exact same input data including rarefaction (Fig. 2i and Fig. 2k unweighted EMP; Fig. 2j and Fig. 2l, weighted EMP; Fig. S3a, b for DartUniFrac GWMC; c and d for exact UniFrac GWMC). PCoA analysis based on Qiita amplicon studies (279,443 samples) ^19^ showed similar results (Fig. S4a and c for unweighted; Fig. S4b and d for weighted). Similar results also were found for large-scale animal gut microbiome datasets and human gut metagenomic datasets (Fig. S5a, b, c and d; Fig. S6a, b, c and d). Procrustes analysis showed that DartUniFrac aligns almost perfectly with exact UniFrac for GWMC dataset (Fig. 1h for unweighted; Fig. S7 for weighted) (M^2^=0.0042 and 0.0039, *P*<0.001) and EMP dataset (Fig. S8). Mantel correlation between DartUniFrac and exact UniFrac for all the datasets agrees with Procrustes analysis (Fig. 1f and g, Mantel r >= 0.98, *P*<0.001; Table S6). Efficient Rejection Sampling (ERS) showed similar PCoA results for EMP dataset (Fig. S9a and b) and Mantel results (Table S7) with exact UniFrac but is ∼60 times slower than DartMinHash (Fig. 2b and Table S8) for the sparse datasets EMP & AGP (Fig. S10b). However, for a denser dataset GWMC with an average sparsity of ∼10.3% (Fig. S10a), ERS is ∼2-3 times faster than DartMinHash (Table S9). To test if DartUniFrac is sensitive to tree size and number of samples, we showed that for a fixed number of samples and sparsity, running time to extract branches via balanced parentheses representation is constant with respect to the number of taxa in the tree (Fig. 2c), consistent with the claim above that BP can scale to billions of taxa. As sparsity increases without changing the number of samples, running time increases proportionally (Fig. 2c). In practice, increasing the number of samples will both increase the number of taxa and total branches. Therefore, DartMinHash sketching time increases proportionally with the number of samples but is generally fast with multi-threaded implementation (Fig. 2d).

To further speed up downstream analysis for a large number of samples (large distance matrix output), we developed a novel a fast Principal Coordinate Analysis (fPCoA) algorithm. The key step is to replace the exact but much slower singular value decomposition (SVD) step with a faster, approximate SVD-- randomized SVD--with controllable error. Among several variants of randomized SVD, we rely on the subspace iteration-type randomized SVD ^20^ as subspace iteration was shown to have fewer vectors to orthogonalize (See Online methods). fPCoA is >100 times faster than exact PCoA and can be ∼25% to ∼33% faster than the fast PCoA in scikit-bio (Table S10). The ordination results showed that fPCoA is nearly identical to exact PCoA for top coordinates (Fig. S11, Procrustes M^2^=0.00001, P<0.001). Because DartUniFrac is much faster, it can now be used to perform statistical resampling tests on UniFrac-based clustering (e.g., UPGMA or neighbor-joining), allowing us to quantify how robust the resulting trees are to sampling noise in community composition, a well-recognized issue in PCR-based amplicon community profiling ^21^. For the GWMC dataset (N = 1,185), UPGMA trees obtained from jackknife resampling with DartUniFrac were highly consistent with those derived from the same jackknife procedure using exact UniFrac distances, indicating similar clustering structure (Fig. S12). For the full EMP & AGP dataset (N = 50,085), 50 rounds of jackknife resampling with DartUniFrac (CPU) completed in under 45 minutes, whereas the same procedure with exact UniFrac (Striped UniFrac) required more than 10 hours (Table S11). With multi-GPU support, it can be finished within 6.3 minutes for DartUniFrac, but it needs 2.1 h with unifrac-binaries (GPU) (Table S11).

The new UniFrac algorithm can greatly accelerate large-scale UniFrac computation for many applications. For example, in the web-enabled microbiome meta-analysis platform Qiita ^19^, users have deposited more than half a million microbiome samples (including metagenomic datasets) and the total number of unique taxa markers is in the tens of millions (including both amplicon sequencing and metagenomic sequencing) (accessed November 2025). DartUniFrac also enables routine cross-study meta-analysis at repository scale and makes sensitivity analyses (e.g., jackknifing, bootstrapping, permutation-based significance testing) computationally feasible. When a fixed reference phylogeny is used—as is typical for close-reference community profiling—new samples can be appended by computing only the new rows/columns against the existing sketches, avoiding re-computation of the full DartUniFrac distance matrix. Together, these capabilities lower the computational cost and energy barrier for million-sample phylogenetic beta-diversity. The new algorithm can only compute unweighted and weighted UniFrac, but not generalized UniFrac ^22^ or variance adjusted UniFrac ^23^ as the latter two UniFrac algorithms cannot be formulated as weighted Jaccard similarity over tree branches. Because of this, both generalized UniFrac and variance adjusted UniFrac remain difficult to run beyond a few thousand samples or more in practice despite their potential advantages in handling abundance-dependent effects, improving power for shifts in rare to moderately abundant taxa and stabilizing noisy abundance differences through variance weighting, respectively. For Earth-Mover Distance UniFrac (EMDUniFrac) ^24, 25^, which was proposed recently and is essentially the numerator of Weighted UniFrac (Fig. S1b), however, we cannot efficiently approximate it via unbiased locality sensitive hashing algorithms ^26^ whereas Weighted MinHash (DartMinHash) for the original UniFrac (both unweighted and weighted) is unbiased. Interestingly, the DartUniFrac algorithm can be equally applied to absolute abundance UniFrac ^27^, where we take cell absolute counts of a given species or strain instead of relative abundance as inputs to build the sample set vectors.

Computational complexity analysis showed that DartUniFrac computing time is dominated by the number of samples, but not the number of taxa while exact UniFrac algorithms are dominated by O(N^2^*T) ^3^, which becomes impractical when the taxa count T increases with sample size—often approximately proportional to N as additional samples reveal new taxa/strains—so the worst runtime can approach O(N^3^) (or even worst) in practice. In fact, none of the packages providing current exact UniFrac implementations supports more than 20 million taxa and 500,000 samples (in part due to input file format limitations), which makes DartUniFrac the only option without additional software engineering effort. Crucially, this scalability is not only about peak throughput but also about feasibility: without sketching, the memory footprint of O(N^2^*T) methods become prohibitive long before computation finishes. By decoupling compute from the raw taxa count and operating on fixed-length signatures, DartUniFrac remains tractable as reference phylogenies and feature catalogs continue to expand (e.g., strain-resolved and spatially resolved metagenomics). DartUniFrac can handle real-world large-scale datasets with different sparsity, e.g., per-sample amplicon denoising ^28^ and pool-sample amplicon denoising ^29, 30^ methods can have different sparsity (∼0.01% and ∼5%, respectively) ^4, 18^. The ERS algorithm is especially efficient relative to DartMinHash for datasets with >4% sparsity (Fig. 2b). With the recent advancement in spatial metagenomics technology at fine resolution ^31^, it is not uncommon to obtain even denser datasets with millions or billions of taxa, especially for homogeneous environments, such as marine and freshwater. The balanced parentheses data structure for representing trees is theoretically optimal ^32^ and can scale to trees with billions of taxa. Both DartMinHash and ERS match the current theoretical lower bound for unbiased estimation of weighted Jaccard similarity for sparse and dense datasets, achieving better accuracy and running speed than BagMinHash and Improved Consistent Weighted Sampling (or ICWS) ^33-35^. Because these estimators recover similarity purely from hash collision counts, the problem of large-scale UniFrac computation in DartUniFrac essentially reduces to computing integer Hamming similarities (normalized) between fixed-length hash signatures. However, integer Hamming similarity between fixed-length integer hash signatures is pure memory-bandwidth-bound, but not compute-bound, indicating that hardware with better memory bandwidth is the most efficient way to speed up DartUniFrac computation. This is consistent with our experiments that DartUniFrac on commodity GPUs (e.g., NVIDIA RTX 6000 Pro and A100) is much faster than on CPUs for the integer Hamming similarity step.

The accuracy of DartUniFrac can be further improved by increasing the sketch size, at additional computational cost—for example, performing integer Hamming similarity on longer vectors/sketches—which in turn places even greater demands on hardware memory bandwidth (both CPU and GPU) to achieve high speed. We also provided both CPU and GPU streaming mode so that we can compute a small portion of the large distance matrix (N^2^ entries) to avoid large RAM requirements for millions of samples. In addition, the main computational bottleneck—the pairwise integer Hamming similarity between sketch vectors—is embarrassingly parallel and can be distributed across many computing nodes, with or without GPUs. The pairwise sketch-level computations decompose naturally into independent subproblems (blocks of the distance matrix) that can be assigned to different nodes with minimal communication overhead.

In downstream analysis, e.g., PCoA, for the accuracy levels relevant to our applications, we find that power iteration-based randomized SVD achieves the desired precision while operating on a smaller working subspace than a randomized block Krylov implementation ^20^. This reduces orthogonalization and memory traffic and yields a modest speedup in practice, consistent with theoretical prediction. More importantly, power iteration naturally supports a streaming/out-of-core implementation: each pass over the distance matrix only requires the current sketch, so the matrix can be processed in blocks. In contrast, randomized block Krylov methods must construct and orthogonalize a full block Krylov basis of dimension, which is difficult to maintain in a streaming setting without storing all intermediate blocks or recomputing them ^36^. For an even larger number of samples (e.g., larger than a million), combined with streaming mode in DartUniFrac, streaming PCoA can reduce memory requirement significantly without sacrificing accuracy.

In summary, DartUniFrac will serve microbiome research for years to come, especially as spatial genomics becomes increasingly popular for microbial communities. It will enable the study of much more diverse environments, such as the soil at fine-grained spatial and temporal resolution and help address microbial ecological and evolutionary questions at unprecedented scales. DartUniFrac paves the way for training deep learning models for microbiomes via providing fast and accurate truth at the millions scale or above.

### Online Content

Any methods, additional references, reporting summaries, source data, extended data, supplementary information, acknowledgements; details of author contributions and competing interests; and statements of data and code availability.

## Methods

### Balanced Parentheses (BP) representation of phylogenetic trees

For a given Newick format tree, we represent the tree using a succinct, bit-level balanced-parentheses encoding instead of a pointer-based tree. During a pre-order traversal, we append an open bit (1) when first visiting a node and a close bit (0) after processing its children, while storing node labels (e.g., taxon IDs) in a compact array in the same order. The resulting bitstring is stored as 64-bit words with a succinct rank/select index and a two-level “pioneer” index for matching parentheses across word boundaries. This allows constant-time navigation operations (first child, next sibling, parent, subtree size, etc.) that touch only a few 64-bit words and small auxiliary arrays, with no pointer chasing (Fig. 1a). The topology is stored in ∼2 bits per parentheses (≈4N bits for N nodes) plus low overhead, so trees with billions of taxa remain memory-feasible (a few GB) and efficiently traversable on modern multi-core CPUs. This balanced parentheses representation has been shown to be theoretically optimal ^32^. See Supplementary Methods for detailed description.

### DartMinHash implementation details

For any given weighted set *X* = {(*i, x*_*i*_)} with non-negative weights (here *i* indicates a branch while *x*_*i*_ is 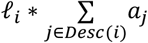, for weighted UniFrac), our bucket-style DartMinHash sketch produces k independent MinHash sketches as follows. First, we perform a parameterization step: choose k and set the expected dart budget *t* ≈ *klnk* + 2*k*. Define a rank threshold *θ* (initially 1.0). Let 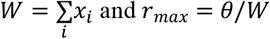 and *r*_*max*_ = *θ*/*W*. Then we perform dyadic tiling of weight–rank space (Fig. S13). For each element (*i, x*_*i*_), we iterate over dyadic scales *v, ρ* ≥ 0. Each pair (*v, ρ*) defines a rectangular “tile” with width Δ*w* = 2^*v*^/*t* ⋅ 2^−*ρ*^ and height Δ*r* = 2^*ρ*^ ⋅ 2^−*v*^. We traverse tiles in increasing w and r, stopping early when *w* ≥ *x*_*i*_ or *r* ≥ *r*_*max*_ (which is 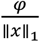, φ controls the upper limit on the rank of darts to return, so that the algorithm returns φ*t* darts in expectation) (Fig. S13). Next, we start a hash-driven Poisson dart count. For every tile, we form a 64-bit key by XORing tabular hashes of *i, v, ρ* and the tile’s local integer offsets. From this key we draw a Poisson(1) count via a CDF (Cumulative Distribution Function) table; this gives the number of darts landing inside the tile. For each dart assigned to a tile, we draw two independent pseudo-random values *U*_*w*_, *U*_*r*_ ∼ *Uniform*(0,1) from the hash and convert them into continuous offsets (w, r) inside the tile: *w* = *w*_0_ + Δ*w* ⋅ *U*_*w*_, *r* = *r*_0_ + Δ*r* ⋅ *U*_*r*_. This yields a point uniformly distributed over the tile area. We accept the dart if *w* < *x*_*i*_ and *r* < *r*_*max*_. Each accepted dart carries an id *h* (another 64-bit tabular hash of the tile key plus a per-dart index) and its rank *r*. After this, a bucketization step was implemented. We maintain k buckets or partitions, each storing the minimum rank seen so far. A separate 64-bit tabular hasher maps h to a bucket index *j* = *h mod k*. If some buckets remain empty after one pass, we increase *θ* and repeat steps 2 to 5 until all k buckets are filled. For all the hashing-related steps, we use simple tabulation hashing (32- and 64-bit), which indexes small random tables with byte/word positions and XORs the looked-up entries ^37^. It is fast, stateless, and provides strong practical independence when composing multiple keys (ids, scales, and tile offsets) for MinHash-like algorithms. It has been shown that for simple MinHash, tabulation hashing, especially mixed tabulation hashing, yields concentration essentially as good as truly random hashing for bucket/partition hashing ^38^. DartMinHash produces per-bucket minima via an exponential race; once each bucket has a winner, the collision indicators across buckets behave like independent Bernoulli trials with mean equal to weighted Jaccard similarity. Under the hashing assumptions (as good as truly random hashing), DartMinHash has variance and concentration that matches the unweighted case in practice, although mixed tabulation hashing provides even stronger guarantees ^38^. For any two resulting sketches with length k, the hash collision probability of IDs across positions is an unbiased estimator of weighted Jaccard similarity and the variance is J(1 − J)/k. Default sketch size k is 2,048 unless specified otherwise.

### Big-O notation for DartMinHash and B-bit DartMinHash

The DartMinHash sketch step big-O notation is: 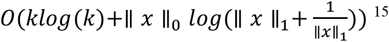, where ∥ *x* ∥_0_ is the number of non-zero elements in a set (one sample), while ∥ *x* ∥_1_ is the sum of weights of all non-zero elements. As long as ∥ *x* ∥_1_ is not extremely small or large, which is the case for real-world microbiome datasets, DartMinHash sketching complexity is simply *O*(*klog*(*k*) + *c* ∗∥ *x* ∥_0_), where c is a small constant. In the case of DartUniFrac, ∥ *x* ∥_0_ will be the average number of active branches for each sample and k is the sketch size. We rewrite the sketching complexity for N samples using T_active_ as ∥ *x* ∥_0_ and S as the sketch size: O(N*T_active_ + N*S*log(S)). Similar to the b-bit MinHash idea ^39^, the 64-bit integer hashes in sketch vectors can be reduced to b=32 or 16 bit (default) by extracting the lower 32 or 16-bit of hashes before computing collision probability ^39^. The noise of the new estimator is 2^−*b*^. For b > 14, the noise can be ignored as 2^−*b*^ is rather small.

### Efficient Rejection Sampling (ERS) implementation details

We first define caps and red–green indexes. For a weighted set *X* = {(*i, x*_*i*_)} (here *i* indicates a branch while *x*_*i*_ is 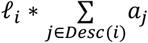, where *a*_*j*_ is the abundance of descendant taxa/leaves under branch *i*) with *x*_*i*_ ≥ 0, choose tight real-valued caps *m*_*i*_ ≥ *max x*_*i*_(*s*) across the dataset and build a cumulative array *cum* with *cum*[0] = 0, *cum*[*i* + 1] = *cum*[*i*] + *m*_*i*_; let 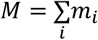. This defines a line [0,M) partitioned into segments [*cum*[*i*], *cum*[*i* + 1]]. A draw *r* ∈ [0, *M*) maps to component i by binary search (‘red–green’ test). Then, for efficiency, we create dense weights for membership tests. Materialize a dense vector w of length D with *w*_*i*_ = *x*_*i*_ (0 otherwise). This allows an O(1) acceptance check once the component index is found. We then generate k independent sequences. For each hash position *j* ∈ {0, …, *k* − 1}, generate a fixed-length sequence *r*_*j*,1_, …, *r*_*j,L*_. Each *r*_*j,t*_ is produced by hashing the key (*j, t*) to a uniform *u* ∈ [0,1) and setting *r* = *Mu*. Next, the red–green acceptance test. Map r to component i by upper bounding in *cum*. The draw is green (accept) if *r* ≤ *cum*[*i*] + *x*_*i*_; otherwise, it is red (reject). Upon first green for position j, emit the pair (id_j_, rank_j_) where *id*_*j*_ is a 64-bit hash derived from the accepted draw r and rank_j_: = *t* (the first-hit index). After the red-green test, a densification (or rotation) was performed. If a position j has no green after L attempts, mark it empty and densify draw a per-j offset *o* ∈ {1, …, *k* − 1} and scan circularly to copy the nearest non-empty bucket’s (id, rank). This procedure is data-independent and preserves the collision-probability estimator. We finally return k pairs (id_j_, rank_j_). For any two sketches, the collision probability of IDs across positions is an unbiased estimator of weighted Jaccard similarity and the variance is J(1 − J)/k. Tight capacities (small M) increase acceptance and allow smaller L. All randomness is via simple tabulation hashing: treat keys as words/bytes, look up in small random tables, and XOR results ^37^. Separate 64/32-bit tabulators drive (i) uniforms *u* for *r*, (ii) stable IDs from *r*, and (iii) densification offsets—yielding fast, stateless, reproducible pseudo randomness suitable for composing keys (*j, t*) and *r*. Let *p* = *Pr*[*green*] =∥ *x* ∥_1_/*M* under caps *m*_*i*_ (with 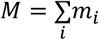). A bucket is empty only if all L trials are rejected, so *p*_*empty*_ = (1 − *p*)^*L*^ ≈ *e*^−*pL*^. We choose L to make empties rare across the sketch, e.g., *kp*_*empty*_ ≤ *ϵ*, giving L≳ *ln*(*k*/*ϵ*)/*p*. Thus, tighter caps (smaller M) and larger sample mass ∥ *x* ∥_1_ allow smaller L; in practice we use a conservative fixed L or adapt L using *p* =∥ *x* ∥_1_/*M*. In microbiome feature tables that are highly sparse (typically 0.1%–10% nonzero), the effective ERS acceptance probability *p* =∥ *x* ∥_1_/*M* is often on the order of 10^−2^ or higher under tight dataset-wide caps; in this regime, L=512 makes empty buckets rare for k=2048 (since *k*(1 −*p*)^*L*^ ≈ *ke*^−*pL*^ ≪ 1). Accordingly, we use L=512 as a conservative default and increase L for datasets where the observed p is smaller. The same b-bit idea can be applied for ERS.

### Sketch cost, size and set dimension

DartUniFrac approximates UniFrac distances by estimating the underlying weighted Jaccard similarity *J*_*w*_ between branch-length–weighted feature vectors using Weighted MinHash (DartMinHash or ERS). Theoretical results for (weighted) MinHash show that the estimator *J*_*w*_ is unbiased and that its variance scales as *Var*(*J*_*w*_) ∝ 1/*k*, where k is the sketch size, and does *not* depend on the ambient dimensionality (for example, the number of taxa or branches in the tree). Intuitively, the sketch samples from the distribution of *nonzero* branch masses in each sample; adding branches with zero mass in both samples does not change this distribution and therefore does not affect the approximation error. Consequently, for a given similarity regime and typical sparsity pattern (i.e., when the number of nonempty taxa per sample remains stable), keeping k fixed yields approximately constant root-mean-square error in *J*_*w*_, and thus in the resulting UniFrac distances, even as the tree grows to include many more taxa and branches. This is the advantage over exact methods, where if samples contain the taxa, these taxa/branches must be built in the stripe in the Striped UniFrac algorithm in a pairwise manner and thus increase the space and computation significantly. For DartUniFrac, due to early stop conditions in DartMinHash, more taxa/branch will only add to the hashing cost per sample, which is constant time operation.

### Implementation, Parallelization, SIMD Hamming and GPU Integer Hamming

DartUniFrac was implemented in pure Rust. All steps including tree traversal, sample vector parsing, DartMinHash/ERS sketching and integer Hamming distance computation are fully parallelized via Rayon library. To further speed up integer Hamming distance computation on CPU, we utilized Simple Instruction Multiple-Data (SIMD) on modern CPUs via Rust standard library portable SIMD API. GPU implementation was based on cudarc (https://crates.io/crates/cudarc), a Rust API around NVIDIA CUDA library, supporting multiple NVIDIA GPUs. Briefly, each sketch vector (representing each sample) was sent from CPU to GPU and each GPU maintains a copy of all sketches. Because the sketch vector is very small (a few kilobytes), GPU memory consumption is rather small even for millions of samples (∼3G with default bbits=16) (Fig. 2h). Each GPU is only responsible for computing a small portion of the integer Hamming similarity computation in a multi-GPU setting. The resulting integer Hamming similarity will be sent to the CPU as long as the computation is finished and stored in CPU RAM. In the streaming settings, a block of the distance matrix will be computed first and then sent to the CPU for writing while the GPU starts computation for next block of the distance matrix. Once the writing of the first block is finished, it will be released from CPU memory (RAM). In practice, writing is faster than GPU computing if block size is not too large compared to the number of samples.

### Fast PCoA based on randomized singular value decomposition (rSVD)

Instead of exact singular value decomposition (SVD) in exact PCoA (the speed limiting step) ^40^, we use a randomized SVD based on randomized subspace iteration (simultaneous power iteration) for datasets with >5,000 samples, rather than the randomized block Krylov method used in scikit-bio ^40^. Subspace iteration typically requires orthogonalizing fewer basis vectors and can be more GEMM-heavy (and thus more cache-friendly for level-3 BLAS), whereas block Krylov methods offer stronger accuracy guarantees per pass ^36^. We perform the required vector orthogonalization using a LAPACK backend via the Rust lax crate. In our regime—dense N×N distance matrices with up to hundreds of thousands of samples and a modest target rank—randomized subspace iteration and randomized block Krylov have essentially the same dominant cost, since both are driven primarily by a small number of passes over the distance matrix (matrix–block multiplies) ^36^. Therefore, only modest speedups are expected from switching between these two approaches for in-memory implementations. The output from fPCoA can be imported to scikit-bio and QIIME 2 directly for visualization. To compare with scikit-bio (v0.7.1) fast PCoA, the ‘fsvd’ option was used to perform randomized block Krylov SVD. K=10 was used for both fast PCoA in scikit-bio and DartUniFrac.

### Benchmarking software packages and datasets generation

We use the original Striped UniFrac ^7^ and hardware optimized Striped UniFrac called unifrac-binaries ^9^ as the truth. For GPU-based comparisons, unifrac-binaries GPU implementation was used. The latter represents the current fastest exact UniFrac implementation with hardware optimization. DartUniFrac v0.3.0 was used for all the benchmarks. EMP data was downloaded from EMP project ^4^. The same processing pipeline was used to obtain OTU feature tables and phylogenetic trees. Briefly, sequence data were error-filtered and trimmed to the length of the shortest sequencing run (90 bp) using the Deblur ^28^. Deblur observation tables were filtered to keep only tag sequences with at least 25 reads total over all samples. For comparison to existing OTU tables, traditional closed-reference OTU picking was done against 16S rRNA databases Greengenes 13.8 ^41^ and SILVA 123 ^42^ using SortMeRNA ^43^, and subsampled open-reference OTU picking was done against Greengenes 13.8. The American Gut Project data was also obtained from original paper ^5^ and a similar processing approach was used to obtain OTU feature table and phylogenetic tree. Then, OTU feature table from EMP and AGP were merged (the same closed reference 16S rRNA gene database and phylogeny were used) to build a combined EMP & AGP dataset. The GMTOL dataset, which is a subset of EMP with samples that were taken from animal gut, was generated by mapping the metadata from EMP. We then also retrieved all amplicon studies (278,499 samples, ∼1 million taxa) from Qiita ^19^ via rediom ^44^ and applied a similar sequence processing pipeline as above to obtain OTU feature tables and phylogeny, but against the most recent Greengenes 2 database ^45^, a comprehensive 16S rRNA-based taxonomy/phylogeny framework anchored in a whole-genome phylogeny. The three-country THDMI (the healthy diet and microbiome initiative) dataset was downloaded from the original paper ^46^. Briefly, quality-controlled reads from shotgun metagenomic sequencing were mapped to Web of Life version 2 ^47^ using Bowtie2 ^48^ with the SHOGUN ^49^ parameter set in paired end mode with a feature table estimated from Woltka ^6^. Then, the abundance of each genome was obtained by counting the number of reads mapped to the genome with a minimum coverage breadth filter (0.1%) ^50^. The phylogenetic tree was provided by the above-mentioned genome database. For simulated datasets, e.g., 100,000, 300,000, 500,000 and 1,000,000 samples, all the taxa from EMP & AGP (875,222 taxa) and Greengenes 2 (23,450,268 taxa) were used as input features and then normal distribution of taxa abundance for each sample was generated with adjustable sparsity (e.g., 0.05%∼5%). The code repository for simulating feature tables can be found in the code availability section.

## Data availability

All the data mentioned are publicly available as mentioned above.

## Code availability

DartUniFrac implementation can be found in GitHub repository: https://github.com/jianshu93/dartunifrac or via Zenodo ^51^. Scripts for reproducing the main figures/results can be found in the scripts folder of the GitHub repository. DartUniFrac (including GPU version) can be easily installed via Bioconda: https://anaconda.org/bioconda/dartunifrac. GitHub repositories of key libraries, including DartMinHash/ERS, succparen, SIMD-Hamming and fPCoA can also be found in the above main DartUniFrac repository. GPU implementation can be found here: https://github.com/jianshu93/DartUniFrac/tree/DartUniFrac-GPU. Code for simulating large numbers of microbiome feature tables can be found here: https://github.com/jianshu93/sparse_features.

## Acknowledgement

We want to thank Tobias Christiani and Otmar Ertl for discussions on the DartMinHash algorithm, and Xiaoyun Li for discussions on implementation details of the Efficient Rejection Sampling algorithm.

## Author contributions

J.Z. designed the algorithm, conducted the proof and implemented the code. M. E. L. and C.L. helped with the mathematical proof. J.Z. did the software benchmarking. D.M. I.S., L.P., Y.W., L.K., S.D. and A.G. helped with the software benchmark. I.S. helped with GPU code compiling. J.Z. wrote the manuscript with inputs from all authors. R.K. supervised the work.

