## Supplemental_files for "Enabling Megascale Microbiome Analysis with DartUniFrac"

#### Supplementary Tables

**Table S1**. Running time for DartUniFrac-CPU and unifrac-binaries-CPU for a million samples, in-memory or streaming mode. Only compute time is considered for in-memory mode without IO. Peak memory was recorded via psrecord for computing the upper triangle of the distance matrix before writing the full DM. The phylogenetic tree with 87,522 taxa was used. All benchmarks were performed on a 64-thread Intel(R) Xeon(R) 6737P CPU with 3T of RAM.

| | **Number of Samples** | Compute time | **Peak Memory (RAM)** | **Mode** |
|---|---|---|---|---|
| DartUniFrac-CPU | 1,000,000 | 1.8 h | 2.3T | In-memory |
| DartUniFrac-CPU | 1,000,000 | 4.5 h | - | Streaming |
| unifrac-binaries-CPU[1] | 1,000,000 | > 20 days | - | In-memory |

[1]Substeps option was used for unifrac-binaries-CPU, which means a small number of pairs were computed at a time to reduce memory consumption. We cannot run a million samples without the --substeps option. Peak memory depends on the number of substeps, default value is 96. To run all samples at the same time, peak memory requirement is larger than > 5T for unifrac-binaries.

**Table S2.** Running time and IO time for GPU based DartUniFrac (in-memory) and unifrac-binaries. Sparsity for the input feature table is 0.1%. For in-memory mode, only the upper triangle of the full matrix (N^2/2) was computed. The phylogenetic tree with 87,522 taxa was used. IO time was dominated by writing the distance matrix.

| GPU* | Memory Bandwidth | Number of Samples | Wall Time | IO time | Algorithm |
|---|---|---|---|---|---|
| RTX6000 Pro (8X)[1] | 1,792 G/s | 500,000 | 4.6 min | 13.1 min | DartUniFrac (In-memory) |
| RTX6000 Pro (4X) | 1,792 G/s | 500,000 | 7.3 min | 13.1 min | DartUniFrac (In-memory) |
| A100 (HMBe) (8X) | 1,935 G/s | 500,000 | 3.0 min | 13.1 min | DartUniFrac (In-memory) |
| RTX6000 Pro (8X) | 1,792 G/s | 300,000 | 2.2 min | 4.5 min | DartUniFrac (In-memory) |
| RTX6000 Pro (4X) | 1,792 G/s | 300,000 | 3.2 min | 4.5 min | DartUniFrac (In-memory) |
| RTX6000 (Ada)(4X) | 960 G/s | 300,000 | 6.6 min | 4.5 min | DartUniFrac (In-memory) |
| L40s (4X) | 864 G/s | 300,000 | 4.0 min | 4.5 min | DartUniFrac (In-memory) |
| A100 (HMBe) (4X) | 1,935 G/s | 300,000 | 4.4 min | 4.5 min | DartUniFrac (In-memory) |
| RTX6000 (Ada) (4X) | 960 G/s | 100,000 | 1.24 min | 0.5 min | DartUniFrac (In-memory) |
| RTX6000 Pro (4X) | 1,792 G/s | 100,000 | 0.55 min | 0.5 min | DartUniFrac (In-memory) |
| L40s (4X) | 864 G/s | 100,000 | 0.76 min | 0.5 min | DartUniFrac (In-memory) |
| A100 (HBM2e) (4X) | 1,935 G/s | 100,000 | 0.82 min | 0.5 min | DartUniFrac (In-memory) |
| A100 (HMBe) | 1,935 G/s | 500,000 | 2931.4 | 13.1 min | unifrac-binaries (substeps) [2] |
| RTX6000 Pro | 1,792 G/s | 500,000 | 2231.4 min | 13.1 min | unifrac-binaries (substeps) |
| RTX6000 Pro | 1,792 G/s | 300,000 | 911.3 min | 4.5 min | unifrac-binaries (substeps) |
| RTX6000 Pro | 1,792 G/s | 100,000 | 43.7 min | 0.5 min | unifrac-binaries (in-mem) |
| RTX6000 (Ada) | 960 G/s | 100,000 | 102.3 min | 0.5 min | unifrac-binaries (in-mem) |
| L40s | 864 G/s | 100,000 | 60.4 min | 0.5 min | unifrac-binaries (in-mem) |
| A100 (HBM2e) | 1,935 G/s | 100,000 | 80.2 min | 0.5 min | unifrac-binaries (in-mem) |

*CPU is a 64-thread Intel(R) Xeon(R) 6737P

[1]We can only run RTX6000 Pro for a maximum 500,000 samples in the in-memory mode because DM (fp32) requires ~1T CPU memory. Nodes supporting other GPUs have less than 1T of memory. We cannot run 1M samples for the in-memory mode as memory requirement is > 4T.

[2]unifrac-binaries (v1.6 through Bioconda, retrieved 10.21.2025) requires >100G GPU memory or more for 300,000 or 500,000 samples. Therefore, the substeps option (96 substeps) option was used to run on > 300k samples. Unifrac-binaries does not support multi-GPU despite 8 GPUs were provided.

**Table S3.** Running time and IO time for GPU based DartUniFrac (in-memory) and unifrac-binaries (partial mode). Sparsity for the input feature table is 0.1%. Only the upper triangle of the full matrix (N^2/2) was computed. The phylogenetic tree with 23,450,268 taxa was used. IO time was dominated by writing the distance matrix.

| GPU* | Memory Bandwidth | Number of Samples | Compute time | IO time | Algorithm |
|---|---|---|---|---|---|
| RTX6000 Pro (8X)[1] | 1,792 G/s | 500,000 | 11.4 min | 13.1 min | DartUniFrac (In-memory) |
| A100(HBM2e) (8X) | 1,935 G/s | 500,000 | 9.0 min | 13.1 min | DartUniFrac (In-memory) |
| A100(HBM2e) (4X) | 1,935 G/s | 500,000 | 11.7 min | 13.1 min | DartUniFrac (In-memory) |
| A100(HBM2e) (2X) | 1,935 G/s | 500,000 | 13.8 min | 13.1 min | DartUniFrac (In-memory) |
| A100(HBM2e) | 1,935 G/s | 500,000 | - | - | unifrac-binaries-GPU (partial mode)[1] |

*CPU is a 128-thread Intel(R) Xeon(R) 6737P

[1]We cannot run unifrac-binaries for 23 million taxa with 0.1% sparsity as the total number of entries goes beyond the default supported number ($2^{32}$).

**Table S4.** Running time for GPU-based DartUniFrac (streaming mode). Sparsity for the input feature table is 0.1%. Note that in streaming mode, the full matrix (N^2) was computed, and it can be 2 times slower than the in-memory mode. The phylogenetic tree with 87,522 taxa was used.

| GPU* | Memory Bandwidth | Number of Samples | Compute time | IO time | Algorithm |
|---|---|---|---|---|---|
| RTX6000 Pro (8X)[1] | 1,792 G/s | 1,000,000 | 21.1 min | 55.3 min | DartUniFrac (Streaming) |
| A100(HBM2e) (8X) | 1,935 G/s | 1,000,000 | 18.4 min | 55.3 min | DartUniFrac (Streaming) |
| RTX6000 Pro (4X) | 1,792 G/s | 1,000,000 | 38.1 min | 55.3 min | DartUniFrac (Streaming) |
| RTX6000 Pro (8X) | 1,792 G/s | 500,000 | 5.5 min | 13.1 min | DartUniFrac (Streaming) |
| RTX6000 Pro (4X) | 1,792 G/s | 500,000 | 11.2 min | 13.1 min | DartUniFrac (Streaming) |
| RTX6000 Pro (8X) | 1,792 G/s | 300,000 | 2.8 min | 4.5 min | DartUniFrac (Streaming) |
| RTX6000 Pro (4X) | 1,792 G/s | 300,000 | 6.6 min | 4.5 min | DartUniFrac (Streaming) |
| RTX6000 Pro | 1,792 G/s | 1,000,000 | > 7 days | 55.3 min | unifrac-binaries-GPU (partial mode)[1] |

*CPU is a 64-thread Intel(R) Xeon(R) 6737P

[1]To run unifrac-binaries for 1 million samples, sparsity was reduced to 0.03% (it can only support up to $2^{32}$ entries), therefore, if the same sparsity (0.1%) was used for unifrac-binaries, it will take approximately 2 times the current running time according to big-O notation.

**Table S5.** Runtime comparison of GPU-based DartUniFrac (in-memory) and unifrac-binaries (GPU) on a commodity GPU (Nvidia GeForce RTX 5090) for N = 50,085 samples from the EMP & AG dataset. A smaller phylogenetic tree was used for this benchmark, containing 307,055 taxa.

| **GPU*** | **Memory Bandwidth** | **Number of Samples** | Wall time | **Algorithm** |
|---|---|---|---|---|
| RTX5090 (1X) | 1,700 G/s | 50,085 | 0.27 min | DartUniFrac (In-memory) |
| RTX5090 (1X) | 1,700 G/s | 50,085 | 3.50 min | unifrac-binaries[2] |

*CPU is a 16-thread AMD Ryzen 9800X3D

[2]unifrac-binaries consumes 10G+ GPU memory for this dataset. The RTX5090 GPU has 32G memory.

**Table S6**. Mantel correlation between DartUniFrac (DartMinHash) and truth obtained from original UniFrac for 5 different datasets.

| **Dataset** | **Number of Samples** | **Unweighted** | | **Weighted** | |
|---|---|---|---|---|---|
| | | **Mantel r** | ***P*** | **Mantel r** | ***P*** |
| GWMC | 1,185 | 0.9967 | <0.001 | 0.9965 | <0.001 |
| GMTOL | 3,964 | 0.9979 | <0.001 | 0.9972 | <0.001 |
| EMP | 27,751 | 0.9954 | <0.001 | 0.9933 | <0.001 |
| AMPLICON-Qiita | 278,499 | 0.9999 | <0.001 | 0.9989 | <0.001 |
| AGP-EMP | 50,085 | 0.99999 | <0.001 | 0.99999 | <0.001 |

**Table S7.** Mantel correlation between DartUniFrac (Efficient Rejection Sampling) and truth obtained from original UniFrac for 4 different datasets.

| | | **Unweighted** | | **Weighted** | |
|---|---|---|---|---|---|
| **Dataset** | **Number of Samples** | **Mantel r** | ***P*** | **Mantel r** | ***P*** |
| GWMC | 1,185 | 0.9823 | <0.001 | 0.9855 | <0.001 |
| GMTOL | 3,964 | 0.9792 | <0.001 | 0.9809 | <0.001 |
| EMP | 27,751 | 0.9908 | <0.001 | 0.9847 | <0.001 |
| AMPLICON-Qiita | 278,499 | 0.9931 | <0.001 | 0.9920 | <0.001 |
| AGP-EMP | 50,085 | 0.9949 | <0.001 | 0.9902 | <0.001 |

**Table S8**. Sketching time comparisons between DartMinHash and ERS (CPU only). All datasets have 0.1% sparsity by simulation. L=1024 was used for ERS. The phylogenetic tree with 87,522 taxa was used.

| | Number of Samples | | |
|---|---|---|---|
| | 50,000 | 100,000 | 300,000 |
| DartMinHash[1] | 1 s | 4 s | 16s |
| ERS[1] | 1.7 min | 4.94 min | 17.8 min |

[1]A a 64-thread Intel(R) Xeon(R) 6737P CPU

**Table S9**. Sketching time comparisons between DartMinHash and ERS. All datasets have 10.3% sparsity by simulation. L=256 was used for ERS. The phylogenetic tree with 87,522 taxa was used.

| | Number of Samples | | |
|---|---|---|---|
| | 50,000 | 100,000 | 300,000 |
| DartMinHash[1] | 3 s | 9 s | 24s |
| ERS[1] | 1.4 s | 2.7s | 9.3 s |

[1]A a 64-thread Intel(R) Xeon(R) 6737P CPU

**Table S10**. Running time comparisons for PCoA on CPU.

| | Number of Samples | Wall time (CPU)[1] |
|---|---|---|
| Scikit-bio[2] | 50,000 | 13.50 s |
| Scikit-bio | 100,000 | 2.32 min |
| Scikit-bio | 300,000 | 24.71 min |
| Scikit-bio | 500,000 | 79.0 0min |
| DartUniFrac-fpcoa | 50,000 | 22.89 s |
| DartUniFrac-fpcoa | 100,000 | 1.92 min |
| DartUniFrac-fpcoa | 300,000 | 18.20 min |
| DartUniFrac-fpcoa | 500,000 | 53.42 min |
| Exact PCoA | 50,000 | 56.7 min |
| Exact PCoA | 100,000 | 283.24 min |
| Exact PCoA | 300,000 | >10 h |
| Exact PCoA | 500,000 | > 24h |

[1]A 64-thread Intel(R) Xeon(R) 6737P CPU
[2]fsvd option was used. Only CPU was used for both scikit-bio and DartUnifrac because storing DM on GPU requires large amount of GPU memory. Both DartUniFrac and scikit-bio use fp32 for this comparison.

**Table S11**. Running time comparisons for Jackknife analysis for the EMP & AGP dataset.

| | **Iterations** | **Number of Samples** | **Wall time (CPU)[1]** | **Wall time (GPU)[2]** | **Dataset** |
|---|---|---|---|---|---|
| Striped UniFrac | 50 | 50,085 | 1624.5 min | - | EMP & AGP |
| Unifrac-binaries | 50 | 50,085 | 278.4 min | 126.8 min | EMP & AGP |
| DartUniFrac | 50 | 50,085 | 43.4 min | 6.3 min | EMP & AGP |

[1]A a 64-thread Intel(R) Xeon(R) 6737P CPU
[2]8 RTX6000 Pro GPUs were used

### Supplementary Figures

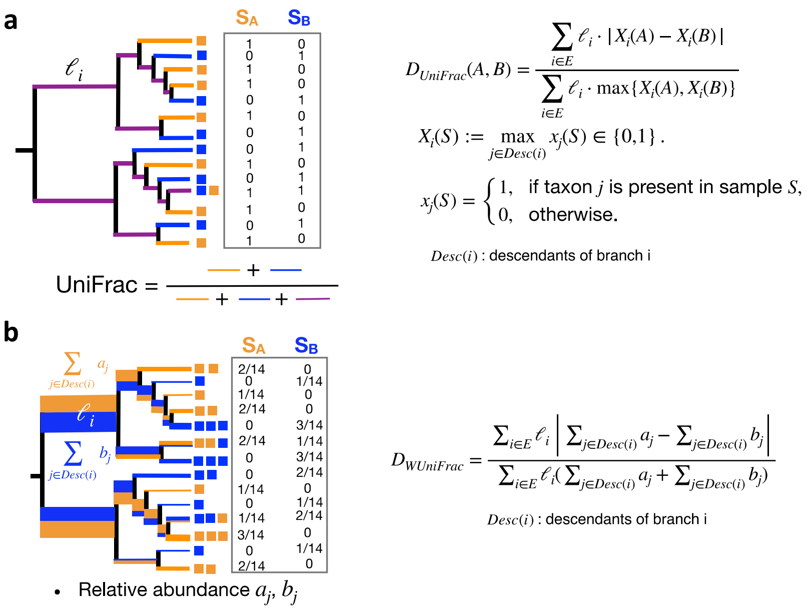


**Figure S1**. Unweighted and Weighted UniFrac in mathematical formula. See Supplementary Methods for detailed explanation of each notation and how both can be related to weighted Jaccard similarity. Note that for Weighted UniFrac, the denominator is different from the original Weighted UniFrac ($D_{WUniFrac} = \frac{\sum_{i \in E} \ell_i \left| \sum_{j \in Desc(i)} a_j - \sum_{j \in Desc(i)} b_j \right|}{\sum_j d_j (a_j + b_j)}$) [1], but it has been proved that they are equivalent [2]. See also Supplementary Methods ("Alternative but equivalent Weighted UniFrac normalization formula") for detailed proof.

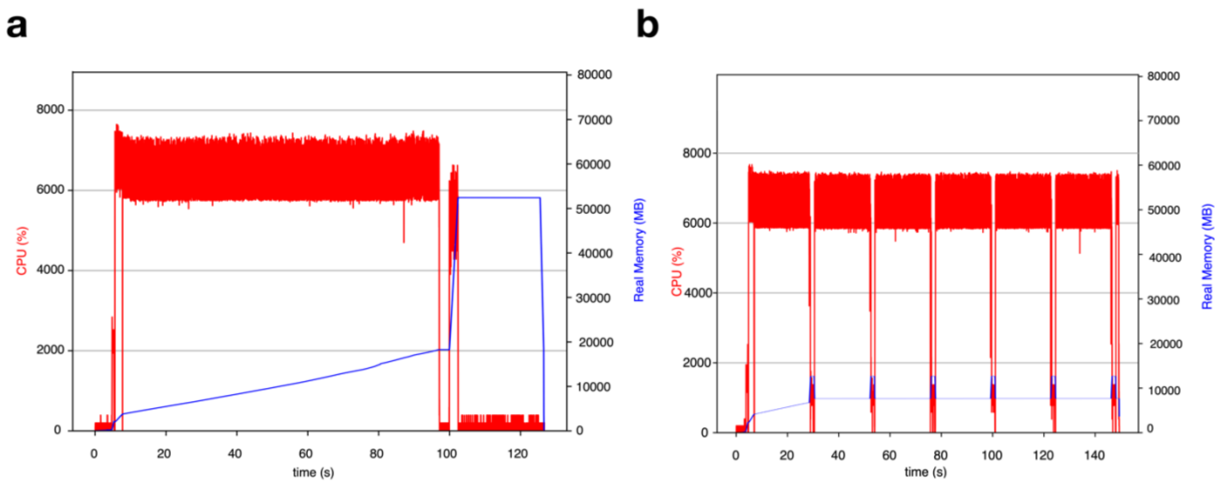


**Figure S2**. CPU and memory usage for DartUniFrac-CPU in in-memory mode (a) and streaming mode (b). Psrecord (https://github.com/astrofrog/psrecord) was used for recording real-time CPU usage and memory consumption. Block size=8192 was used for streaming mode. The EMP & AG dataset was used for this CPU and memory usage analysis.

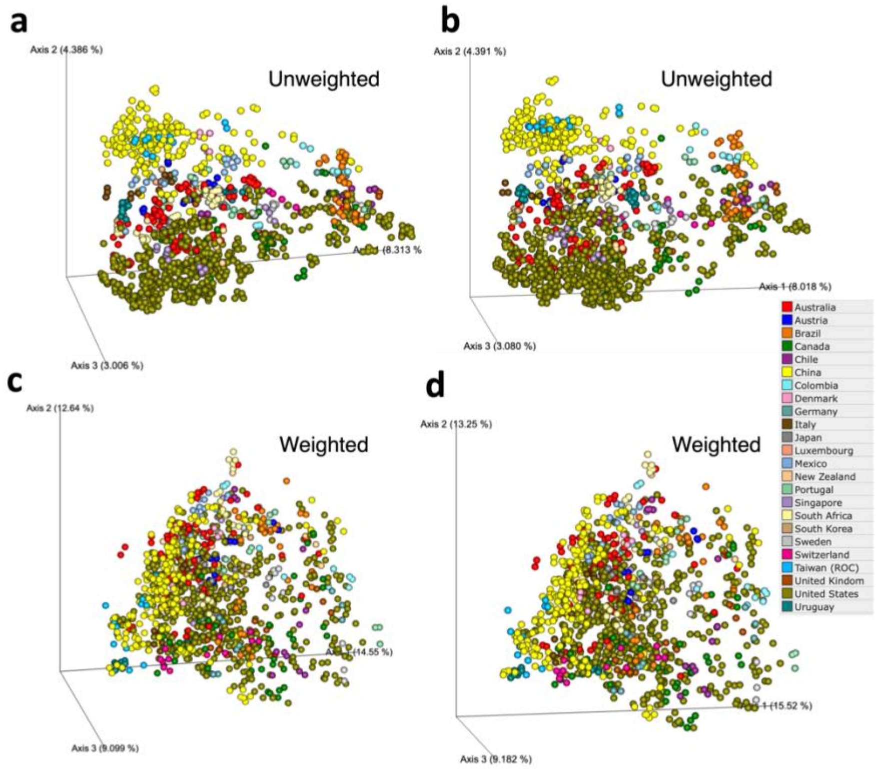


**Figure S3**. PCoA visualization based on DartUniFrac (a and c) and exact UniFrac (b and d) for the GWMC dataset. Samples were colored by country.

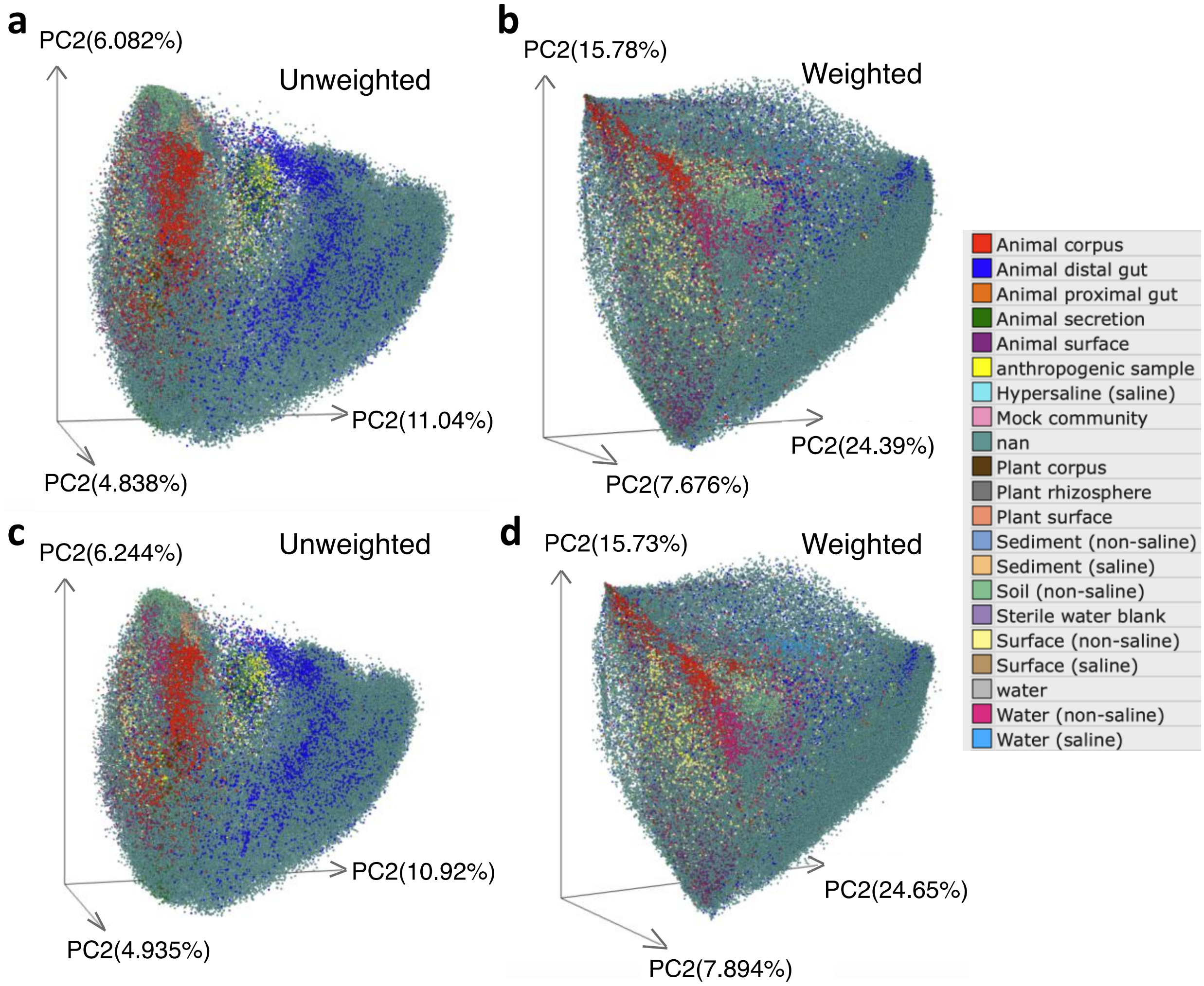


**Figure S4**. PCoA visualization for the Qiita amplicon dataset, unweighted and weighted for exact UniFrac (a and b) and DartUniFrac (c and d). Samples were colored by environmental type.

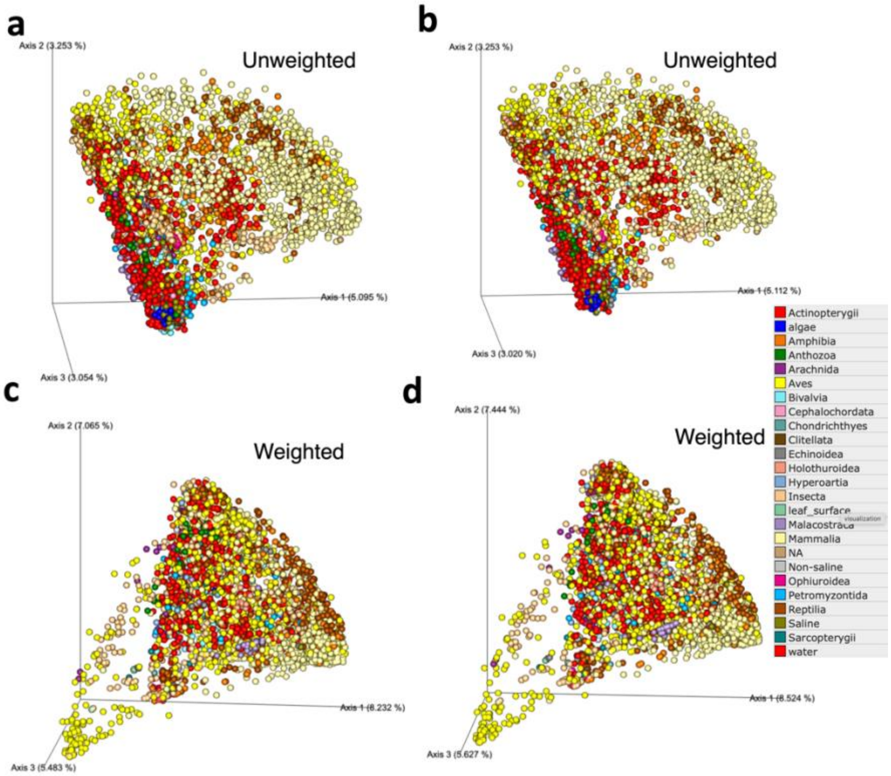


**Figure S5**. PCoA visualization based on DartUniFrac (a and c) and exact UniFrac (b and d) for the animal gut microbiome dataset. Samples were colored by host animal family.

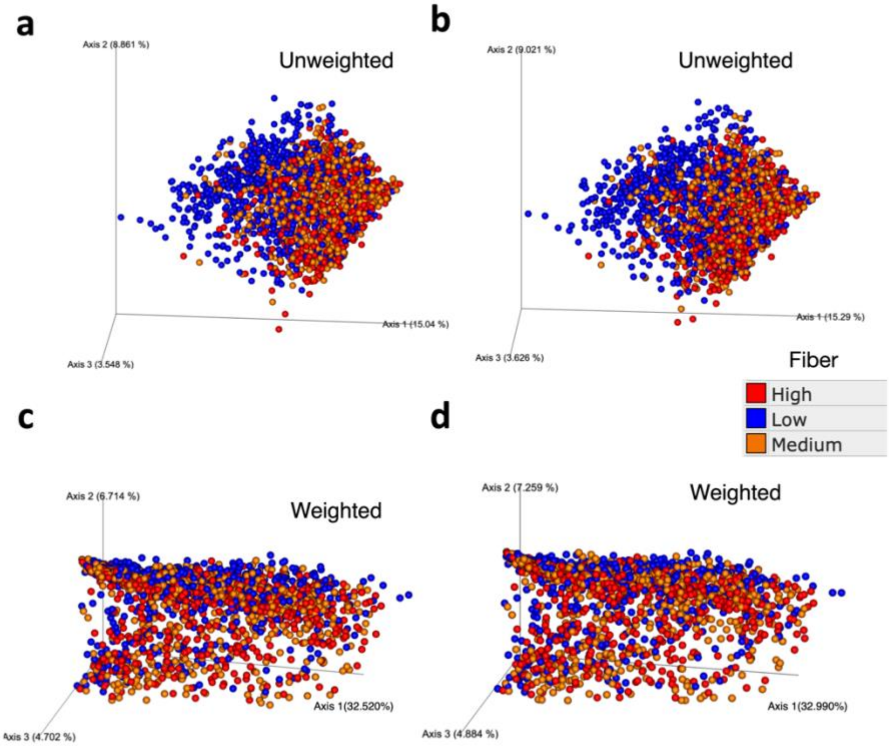


**Figure S6**. PCoA visualization based on DartUniFrac (a and c) and exact UniFrac (b and d) for the human gut metagenomic dataset. Samples were colored by fiber intake. See Online Methods for details on how the data were generated.

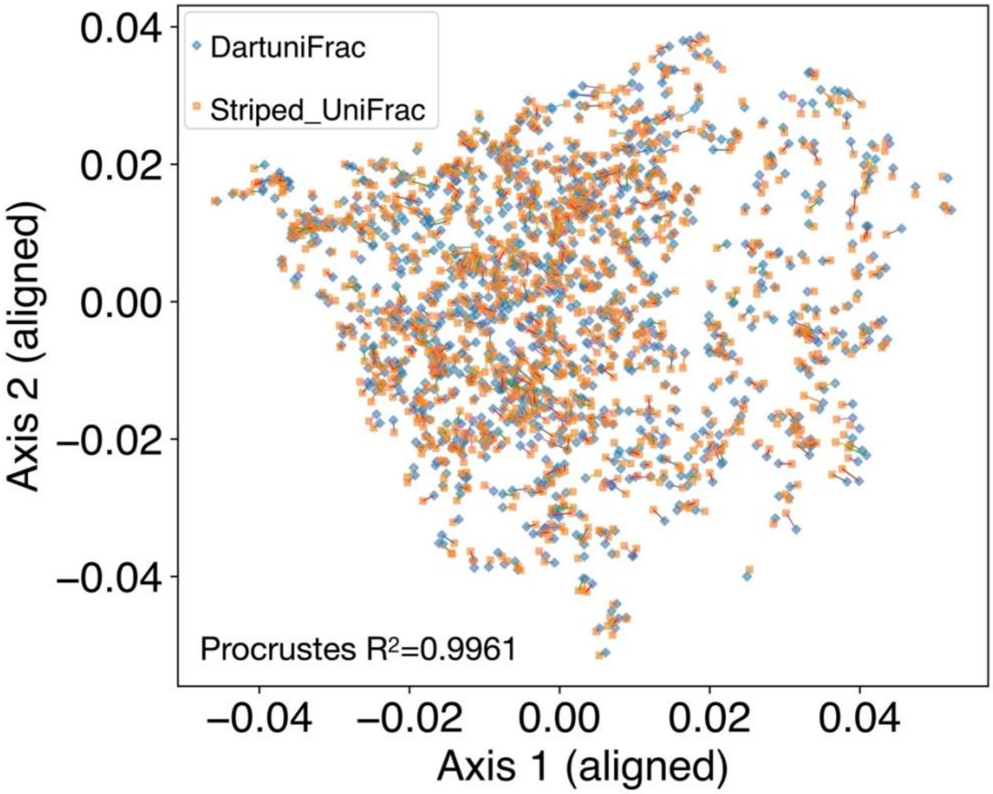


**Figure S7**. Procrustes analysis between DartUniFrac and exact UniFrac for the Global Water Microbiome Consortia (GWMC) dataset (weighted). Procrustes $R^2$ is shown in the plot.

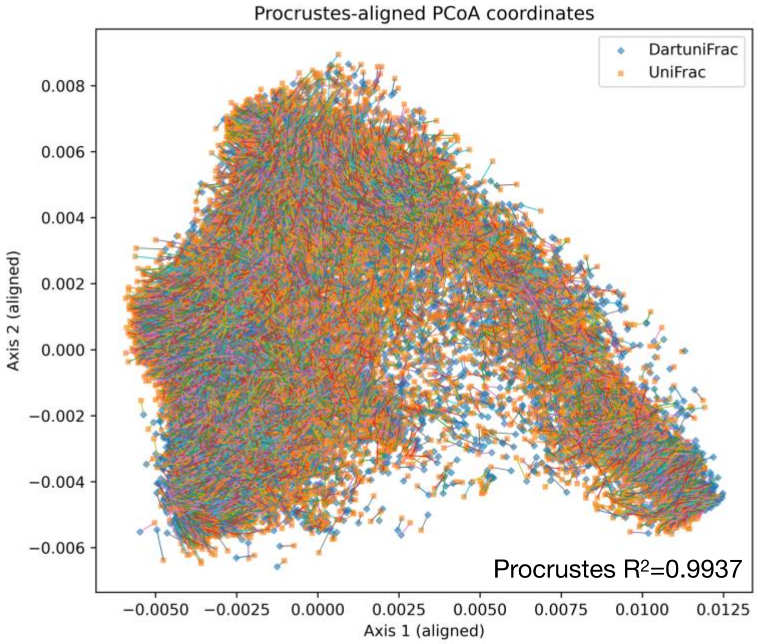


**Figure S8**. Procrustes analysis between DartUniFrac and exact UniFrac for the EMP dataset (weighted). Procrustes $R^2$ is shown in the plot.

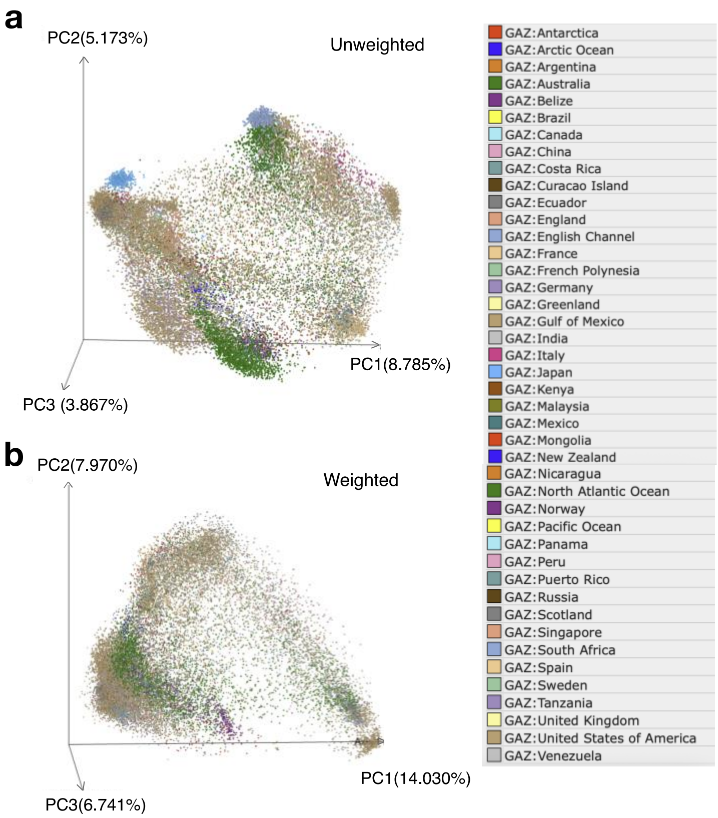


**Figure S9**. PCoA visualization based on DartUniFrac (Efficient Rejection Sampling) for the EMP dataset, unweighted (a) and weighted (b). Samples were colored by country. See **Figure S3** for legend.

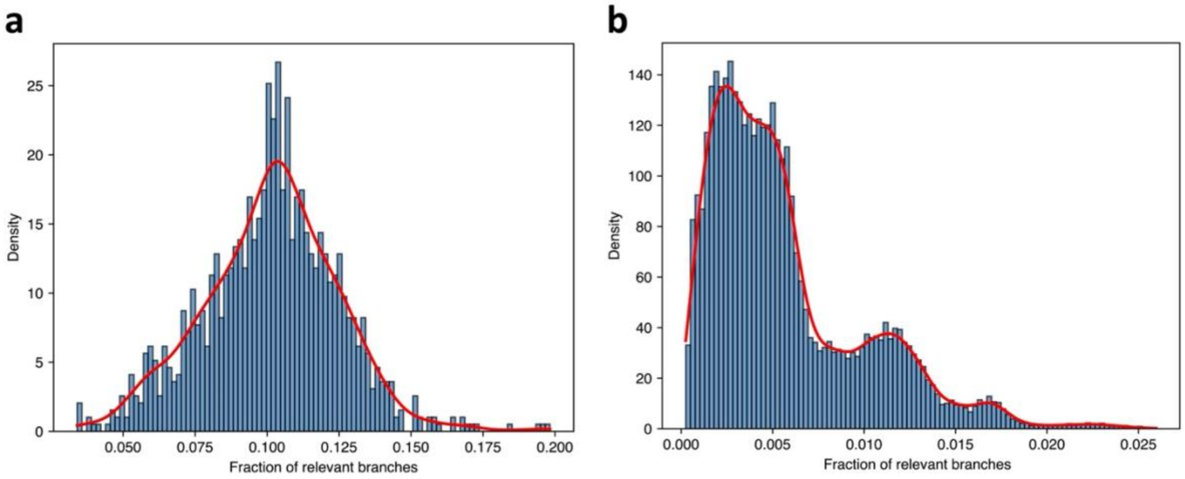


**Figure S10**. Dataset sparsity for GWMC (Global Water Microbiome Consortia, N=1,085) (a) and EMP (Earth Microbiome Project, N=27,000) (b). Sparsity was measured as the fraction of tree branches that will be used as input for sketching vectors out of the total number of branches for each sample (**Figure 1**a). This sparsity definition is related to sample vector but not the same and is more relevant as DartMinHash input is directly related to the number of branches relevant for each sample but not sample vectors.

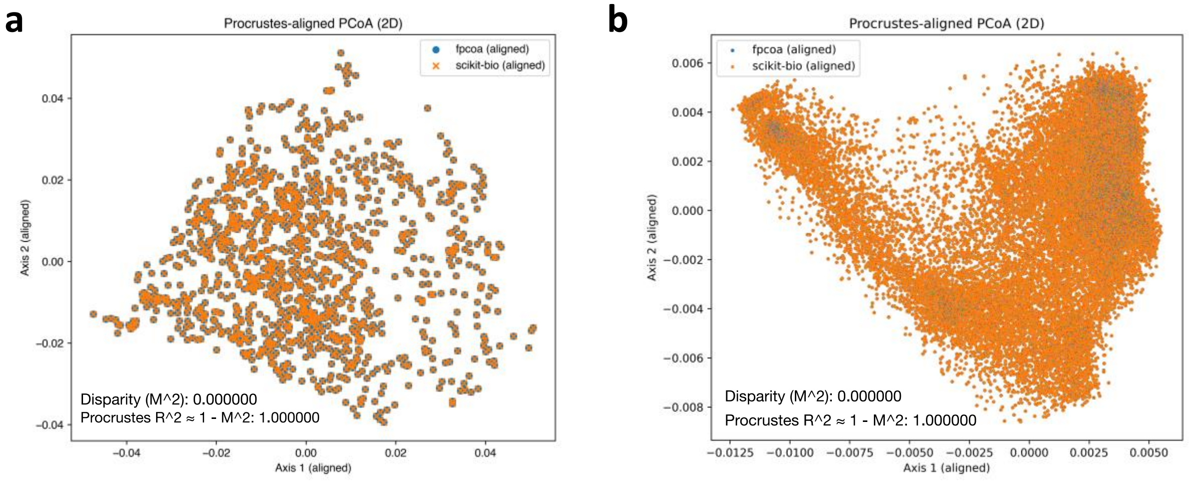


**Figure S11.** Procrustes analysis between fPCoA and scikit-bio exact PCoA for the Global Water Microbiome Consortia (GWMC) dataset (a) and EMP dataset (b). Procrustes $R^2$ is shown in the plot. Exact Weighted UniFrac was used as the input DM for both fpcoa and scikit-bio exact PCoA.

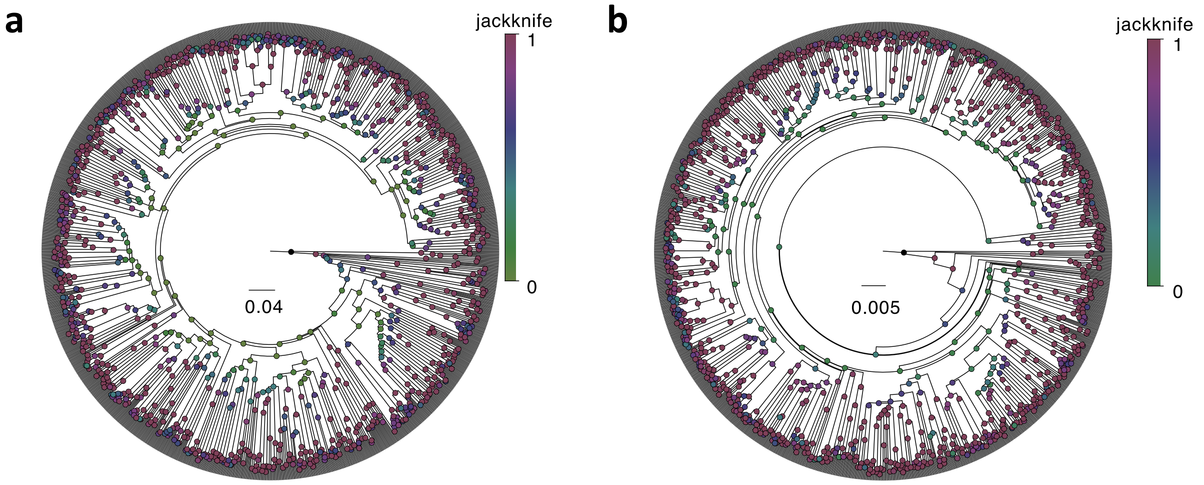


**Figure S12**. Consensus UPGMA tree with Jackknife support for DartUniFrac (a) and exact UniFrac (b). Cophenetic unit-correlation distance: 0.118578 (Pearson r $\approx$ 0.762845) between the 2 UPGMA trees.

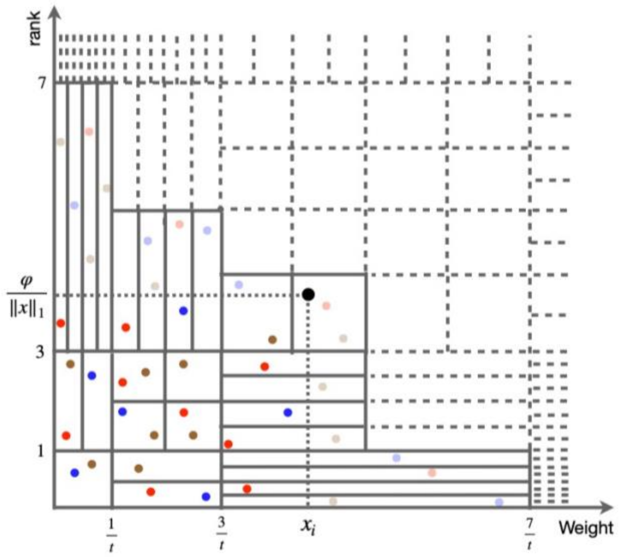


**Figure S13**. Dyadic tiling of weight–rank space for DartMinHash. The weight–rank was discretized into regular rectangles. The number of points generated in each rectangle is Poisson distributed. Points with x larger than the weighted of $x_i$ and y larger than $\frac{\varphi}{\|x\|_1}$, will be rejected. See Online Methods for detailed description of dyadic tiling.

**Supplementary methods**

#### Balanced parentheses Newick representation

For a given Newick format tree, we stored the phylogenetic tree in a succinct, bit-level representation rather than as an explicit pointer-based tree. The topology of the tree is encoded as a balanced parentheses sequence, and node labels (e.g., taxon IDs or internal node identifiers) are stored in a separate, compact array. Let the rooted tree have N nodes (tips + internal nodes). We first parse the Newick tree and traverse it in depth-first pre-order. For each node, when we first visit the node, we append an open parenthesis bit (1) to a global bitstring and append its label to a label array. After we have processed all of its children, we append a close parenthesis bit (0). To simplify indexing, we add a virtual root that wraps the entire tree, so the final bitstring has length 2(N + 1), and each non-virtual node corresponds to one open parenthesis in this sequence. Node labels are stored in the order in which nodes are first visited, so a node's preorder ID is simply its index in the label array. The parentheses bitstring is stored as a contiguous vector of 64-bit machine words. Each word supports constant-time bit operations (bit get/set, population count, rank within the word, and local parenthesis matching) via a small number of arithmetic and bitwise instructions. On top of this, we maintain a succinct index that supports rank and select operations over the whole bitstring: $rank_1(i)$ returns the number of 1 bits in positions 0..i, and $select_1(k)$ returns the position of the k-th 1. These operations are implemented using a multi-level blocking scheme: the bitstring is partitioned into large, middle, and small blocks, and for each block we store only aggregate counts and compact deltas for the positions of 1 bit. This "sparse nearest-neighbor dictionary" over the 1s provides O(1) rank/select in practice with low memory overhead. Tree navigation (child, sibling, parent, subtree size, etc.) is implemented purely in terms of these rank/select and parenthesis-matching operations. A node is represented by the index of its open parenthesis and its preorder label index. The first child of a node at bit position i is found by checking whether bit i+1 is an open parenthesis; if so, the node at i+1 is the first child. The next sibling is found by first locating the matching close parenthesis for position i, then examining the bit immediately after that close: if it is an open parenthesis, it starts the next sibling's subtree. The parent of a node can be obtained by finding the closest unmatched open parenthesis to the left, which again is expressed via rank/select and matching operations. All these operations touch only a handful of 64-bit words and a few small auxiliary arrays, and do not involve pointer chasing or heap allocations. The key non-trivial operation is matching parentheses at large distances given an open parenthesis at position i, find the matching close parenthesis. For pairs that close within the same 64-bit word, we use a lightweight word-local scan. For far pairs (those crossing word boundaries), we build an additional "pioneer" structure. During a single pass over the bitstring we identify far pairs whose open and close parentheses lie in different blocks, mark those positions as pioneers in a second bitstring, and compress this pioneer sequence into its own balanced-parentheses structure. A succinct rank/select index over the pioneer flags lets us map from an arbitrary position in the original sequence to the corresponding location in the pioneer sequence and back. Matching a far parenthesis then proceeds in two steps: (i) locate the nearest pioneer open parenthesis at or before the query position and match it within the pioneer-only sequence, and (ii) refine the match within the local blocks using a small word-level search. This two-level design ensures that parenthesis matching remains effectively constant time even when the enclosing subtree spans many megabytes or gigabytes of bits. Once built, the entire tree topology is represented using approximately 2 bits per parenthesis, i.e., about 4N bits (~0.5 GB for N = 2*10^9 nodes), plus lower order overhead for the rank/select and pioneer indexes. Node labels are stored in a dense array (Vec<L>), which can hold compact integer IDs for leaves and/or internal nodes; labels for internal nodes can be omitted or compressed further if only taxa are needed. Because both the bitstring and label array are contiguous and immutable after construction, they can be shared read-only across many threads or processes (e.g., via reference counting), and each worker only needs to maintain a small node handle (a couple of indices) for navigation. Overall, this balanced-parentheses representation provides a theoretically optimal succinct encoding of rooted ordered trees: it stores the topology in space close to the information-theoretic lower bound while supporting constant-time navigation primitives. In practice, this yields near–constant-time tree navigation with excellent cache locality and a memory footprint small enough to handle trees with on the order of billions of taxa on modern multi-core CPUs.

**Alternative but equivalent Weighted UniFrac normalization formula**

The original normalized weighted UniFrac ( $D_{WUniFrac} = \frac{\sum_{i \in E} \ell_i \left| \sum_{j \in Desc(i)} a_j - \sum_{j \in Desc(i)} b_j \right|}{\sum_j d_j (a_j + b_j)}$ ) defined a denominator of the form: $D_{tips} = \sum_{j \in T} d_j (a_j + b_j)$, where $a_j$, $b_j$ are relative abundances at tip $j$ (taxa $j$) in samples A and B, respectively, while $d_j$ is the total branch length from the root $r$ to taxa $j$(**Figure S1**b). Substitute the definition of $d_j$: $d_j = \sum_{i \in path(r \to j)} \ell_i$, where $\ell_i$ is the length of branch i∈E.

So: $D_{tips} = \sum_{j \in T} \left( \sum_{i \in path(r \to j)} \ell_i \right) (a_j + b_j)$ becomes a double sum. Since the number of items in each sum is finite, we can swap them: $D_{tips} = \sum_{j \in T} \sum_{i \in path(r \to j)} \ell_i (a_j + b_j) = \sum_{i \in E} \ell_i \sum_{j \in T : i \in path(r \to j)} (a_j + b_j)$.

For the inner sum, a node "$j$ such that edge $i$ is on the path from the root to $j$" is exactly a node "$j$ in the subtree below edge $i$" or simply a descendant of edge $i$. So, we've proved: $\sum_j d_j (a_j + b_j) = \sum_i \ell_i ( \sum_{j \in Desc(i)} a_j + \sum_{j \in Desc(i)} b_j )$. Therefore, original Weighted UniFrac in Lozupone et.al., 2007[1]: $D_{WUniFrac} = \frac{\sum_{i \in E} \ell_i \left| \sum_{j \in Desc(i)} a_j - \sum_{j \in Desc(i)} b_j \right|}{\sum_j d_j (a_j + b_j)}$, can be rewritten as: $D_{WUniFrac} = \frac{\sum_{i \in E} \ell_i \left| \sum_{j \in Desc(i)} a_j - \sum_{j \in Desc(i)} b_j \right|}{\sum_{i \in E} \ell_i (\sum_{j \in Desc(i)} a_j + \sum_{j \in Desc(i)} b_j)}$.

1. Lozupone, C.A., Hamady, M., Kelley, S.T. & Knight, R. Quantitative and qualitative β diversity measures lead to different insights into factors that structure microbial communities. *Applied and environmental microbiology* **73**, 1576-1585 (2007).
2. Chang, Q., Luan, Y. & Sun, F. Variance adjusted weighted UniFrac: a powerful beta diversity measure for comparing communities based on phylogeny. *BMC bioinformatics* **12**, 118 (2011).

**Unweighted and Weighted UniFrac are essentially Weighted Jaccard Similarity**

Here we show that:

(a) the *unweighted* UniFrac distance between two samples $A, B$ can be written as

$$D_{\text{UniFrac}}(A, B) = 1 - J_w(\mathbf{x}, \mathbf{y}),$$

where

$$J_w(\mathbf{x}, \mathbf{y}) = \frac{\sum \min\{x_i, y_i\}}{\sum \max\{x_i, y_i\}},$$

i.e., one minus a weighted Jaccard similarity on branch-presence indicators;

(b) the *normalized weighted* UniFrac distance is the monotone transform

$$D_{\text{WUF}}(A, B) = \frac{1 - J_w(\mathbf{x}, \mathbf{y})}{1 + J_w(\mathbf{x}, \mathbf{y})},$$

where $J_w$ is the weighted Jaccard similarity on edge-mass vectors $x_i = \ell_i A_i$ and $y_i = \ell_i B_i$.

### 1 Trees, Samples, and Edge-Level Quantities

Let $T = (V, E)$ be a rooted phylogenetic tree with branch (edge) set $E$ and strictly positive branch lengths

$$\ell_i > 0, \qquad i \in E.$$

Let $L \subseteq V$ be the set of leaves (taxa).

#### 1.1 Presence/absence representation (unweighted case)

For a sample $S$, presence/absence at leaf $j \in L$ is encoded by

$$x_j(S) = \begin{cases} 1, & \text{if taxon } t \text{ is present in } S, \\ 0, & \text{otherwise.} \end{cases}$$

For each edge $i \in E$, let $\text{Desc}(i) \subseteq L$ denote the set of leaves that are descendants of $i$, and define the branch-level presence indicator

$$X_i(S) := \max_{j \in \text{Desc}(i)} x_j(S) \in \{0, 1\}. \tag{1}$$

Thus $X_i(S) = 1$ if and only if at least one descendant taxon under edge $i$ is present in sample $S$.

#### 1.2 Abundance representation (weighted case)

For weighted UniFrac, a sample is represented by nonnegative leaf abundances $\{a_j\}_{j \in L}$ (e.g., relative abundances). For each edge $i \in E$, define the descendant mass

$$A_i := \sum_{j \in \text{Desc}(i)} a_j \quad (\geq 0), \tag{2}$$

and analogously for another sample $B$ with leaf abundances $\{b_j\}_{j \in L}$,

$$B_i := \sum_{j \in \text{Desc}(i)} b_j \quad (\geq 0).$$

We will later collect these into edge-mass vectors $A = (A_i)_{i \in E}$ and $B = (B_i)_{i \in E}$.

#### 2 Weighted Jaccard (Ruzicka) Similarity

Let $d \geq 1$. For nonnegative vectors $\mathbf{u}, \mathbf{v} \in \mathbb{R}^d_+$, define the *weighted Jaccard* (Ruzicka) similarity by

$$J_w(\mathbf{u}, \mathbf{v}) := \frac{\sum_{i=1}^{d} \min\{u_i, v_i\}}{\sum_{i=1}^{d} \max\{u_i, v_i\}}, \tag{3}$$

whenever the denominator is strictly positive.

The associated *weighted Jaccard distance* is

$$d_J(\mathbf{u}, \mathbf{v}) := 1 - J_w(\mathbf{u}, \mathbf{v}),$$

which is known to be a metric (non-negativity, symmetry, and triangle inequality).

#### 3 Unweighted UniFrac as a Weighted Jaccard Distance

The classical unweighted UniFrac distance between samples $A$ and $B$ is defined as the total branch length unique to one sample divided by the total branch length present in at least one sample [1]. In terms of the branch-presence indicators $X_i(\cdot)$ in (1),

$$D_{\text{UniFrac}}(A, B) = \frac{\sum_{i \in E} \ell_i \, |X_i(A) - X_i(B)|}{\sum_{i \in E} \ell_i \, \max\{X_i(A), X_i(B)\}}. \tag{4}$$

**Theorem 1** (Unweighted UniFrac as 1−weighted Jaccard)**.** *Let $X(A) = (X_i(A))_{i \in E}$ and $X(B) = (X_i(B))_{i \in E}$, with weights $w_i := \ell_i$ for $i \in E$. Then*

$$D_{\text{UniFrac}}(A, B) = 1 - \frac{\sum_{i \in E} \ell_i \min\big(X_i(A), X_i(B)\big)}{\sum_{i \in E} \ell_i \max\big(X_i(A), X_i(B)\big)}. \tag{5}$$

*Proof.* For each edge $i \in E$ with indicators $X_i(A), X_i(B) \in \{0, 1\}$, we have

$$|X_i(A) - X_i(B)| = \max\{X_i(A), X_i(B)\} - \min\{X_i(A), X_i(B)\}.$$

Multiplying by $\ell_i$ and summing over $i \in E$ yields

$$\begin{aligned}\sum_{i \in E} \ell_i \, |X_i(A) - X_i(B)| &= \sum_{i \in E} \ell_i \big(\max\{X_i(A), X_i(B)\} - \min\{X_i(A), X_i(B)\}\big) \\ &= \sum_{i \in E} \ell_i \, \max\{X_i(A), X_i(B)\} - \sum_{i \in E} \ell_i \, \min\{X_i(A), X_i(B)\}.\end{aligned}$$

Dividing both sides by $\sum_{i \in E} \ell_i \, \max\{X_i(A), X_i(B)\}$ (assumed $> 0$) gives

$$\frac{\sum_i \ell_i \, |X_i(A) - X_i(B)|}{\sum_i \ell_i \, \max\{X_i(A), X_i(B)\}} = 1 - \frac{\sum_i \ell_i \, \min\{X_i(A), X_i(B)\}}{\sum_i \ell_i \, \max\{X_i(A), X_i(B)\}}.$$

The left-hand side is $D_{\text{UniFrac}}(A, B)$ by (4), and the right-hand side is exactly (5). □

**Corollary 1** (Absorbing branch lengths into $x, y$). *Define edge-weighted vectors*

$$x_i := \ell_i \, X_i(A), \qquad y_i := \ell_i \, X_i(B), \qquad i \in E. \tag{6}$$

*Then*

$$D_{\mathrm{UniFrac}}(A, B) = 1 - J_w(\mathbf{x}, \mathbf{y}) = 1 - \frac{\sum_{i \in E} \min(x_i, y_i)}{\sum_{i \in E} \max(x_i, y_i)}. \tag{7}$$

*Proof.* From (6), we have

$$\min(x_i, y_i) = \ell_i \min\bigl(X_i(A), X_i(B)\bigr), \qquad \max(x_i, y_i) = \ell_i \max\bigl(X_i(A), X_i(B)\bigr),$$

because $\ell_i > 0$ and $X_i(A), X_i(B) \in \{0, 1\}$. Substituting into

$$J_w(\mathbf{x}, \mathbf{y}) = \frac{\sum_i \min(x_i, y_i)}{\sum_i \max(x_i, y_i)}$$

gives

$$J_w(\mathbf{x}, \mathbf{y}) = \frac{\sum_i \ell_i \min(X_i(A), X_i(B))}{\sum_i \ell_i \max(X_i(A), X_i(B))},$$

which coincides with the fraction in (5). Thus (7) follows. □

### 4 Normalized Weighted UniFrac as a Transform of Jaccard

For weighted UniFrac, following [2], the *normalized weighted UniFrac* distance between samples $A$ and $B$ is defined as:

$$D_{\mathrm{WUF}}(A, B) = \frac{\sum_{i \in E} \ell_i \, |A_i - B_i|}{\sum_{i \in E} \ell_i \, (A_i + B_i)}. \tag{8}$$

Introduce the edge-mass vectors $\mathbf{x}, \mathbf{y} \in \mathbb{R}_+^{|E|}$ by

$$x_i := \ell_i A_i, \qquad y_i := \ell_i B_i, \qquad i \in E. \tag{9}$$

Then (8) becomes

$$D_{\mathrm{WUF}}(A, B) = \frac{\sum_i |x_i - y_i|}{\sum_i (x_i + y_i)}, \tag{10}$$

which is the canonical Bray–Curtis form applied to $(\mathbf{x}, \mathbf{y})$.

**Theorem 2** (Normalized weighted UniFrac as a transform of $J_w$). *Let $x, y$ be defined by (9). Then*

$$D_{\mathrm{WUF}}(A, B) = \frac{1 - J_w(\mathbf{x}, \mathbf{y})}{1 + J_w(\mathbf{x}, \mathbf{y})}, \tag{11}$$

*where $J_w$ is the weighted Jaccard similarity $J_w(\mathbf{x}, \mathbf{y}) = \sum_i \min(x_i, y_i) / \sum_i \max(x_i, y_i)$.*

*Proof.* Set

$$X := \sum_i \max\{x_i, y_i\}, \qquad M := \sum_i \min\{x_i, y_i\}.$$

For each coordinate $e$,

$$|x_i - y_i| = \max\{x_i, y_i\} - \min\{x_i, y_i\}, \qquad x_i + y_i = \max\{x_i, y_i\} + \min\{x_i, y_i\}.$$

Summing these identities over $e$ gives

$$\sum_i |x_i - y_i| = X - M, \qquad \sum_i (x_i + y_i) = X + M.$$

Substituting into (10) yields

$$D_{\mathrm{WUF}}(A, B) = \frac{X - M}{X + M}.$$

Dividing numerator and denominator by $X > 0$ gives

$$D_{\mathrm{WUF}}(A, B) = \frac{1 - M/X}{1 + M/X}.$$

But $M/X = J_w(\mathbf{x}, \mathbf{y})$ by definition, hence (11) follows. □

**Corollary 2** (Invertible mapping)**.** *Whenever* $J_w(\mathbf{x}, \mathbf{y}) \in [0, 1]$,

$$J_w(\mathbf{x}, \mathbf{y}) = \frac{1 - D_{\mathrm{WUF}}(A, B)}{1 + D_{\mathrm{WUF}}(A, B)}.$$

*Equivalently, writing* $d_J(\mathbf{x}, \mathbf{y}) := 1 - J_w(\mathbf{x}, \mathbf{y})$*, we have*

$$D_{\mathrm{WUF}}(A, B) = \frac{d_J(\mathbf{x}, \mathbf{y})}{2 - d_J(\mathbf{x}, \mathbf{y})}.$$
